# How a mouse licks a spout: Cortex-dependent corrections as the tongue reaches for, and misses, targets

**DOI:** 10.1101/655852

**Authors:** Tejapratap Bollu, Brendan Ito, Sam C. Whitehead, Brian Kardon, James Redd, Mei Hong Liu, Jesse H. Goldberg

## Abstract

Precise tongue control is necessary for drinking, eating, and vocalizing^1, 2^. Yet because tongue movements are fast and difficult to resolve, neural control of lingual kinematics remains poorly understood. Here we combine kilohertz frame-rate imaging and a deep learning based neural network to resolve 3D tongue kinematics in mice drinking from a water spout. Successful licks required previously unobserved corrective submovements (CSMs) which, like online corrections during primate reaches^3–10^, occurred after the tongue missed unseen, distant, or displaced targets. Photoinhibition of anterolateral motor cortex (ALM) impaired online corrections, resulting in hypometric licks that missed the spout. ALM neural activity reflected upcoming, ongoing, and past CSMs, as well as errors in predicted spout contact. Though less than a tenth of a second in duration, a single mouse lick exhibits hallmarks of online motor control associated with a primate reach, including cortex-dependent corrections after misses.

## Main Text

Accurate goal-directed behavior requires the constant monitoring and correction of ongoing movements. For example, when reaching to an unseen, uncertain or displaced target, errors are estimated and compensated for in real time, resulting in corrective submovements (CSMs) that redirect the hand to its target^3–10^.

Many animals, including humans and rodents, have prehensile tongues that reach out of the oral cavity to contact objects such as food, water and conspecifics^1^. Natural behaviors such as licking, eating, grooming and speaking require fast and accurate tongue movements^1, 2^, yet mechanisms of lingual control remain poorly understood. Even in tractable model systems such as rodents, where licking is used to study movement initiation, planning, and decision making, licks are usually measured as a binary register of whether or not a tongue contacts a spout or transects an IR beam^11–16^ or with 2-dimensional tracking^17, 18^. It remains unclear how a tongue reaches an unseen target such as a water spout.

### High speed tongue imaging identifies corrective submovements important for spout contact

To precisely resolve 3D tongue kinematics, we imaged the tongue at 1 kHz in two planes and trained two deep neural networks^19^ to identify and segment the tongue from side and bottom views (Fig.1a-e, Methods). Using hull reconstruction to build a 3D model of the tongue^20^, we estimated the tongue tip in each frame to achieve millisecond timescale resolution of the lick trajectory (Fig. 1f, Extended Data fig. 1, Movie S1, Methods). Mice were trained to withhold licking for at least one second to earn an auditory cue, and then to lick the spout within 1.3 seconds from the cue to earn a water reward (Fig. 1a, Methods). Cues caused bouts of licking, as previously observed in head-fixed mouse setups where the spout could not be directly seen^11, 17^ (Fig. 1b).

**Fig. 1.**
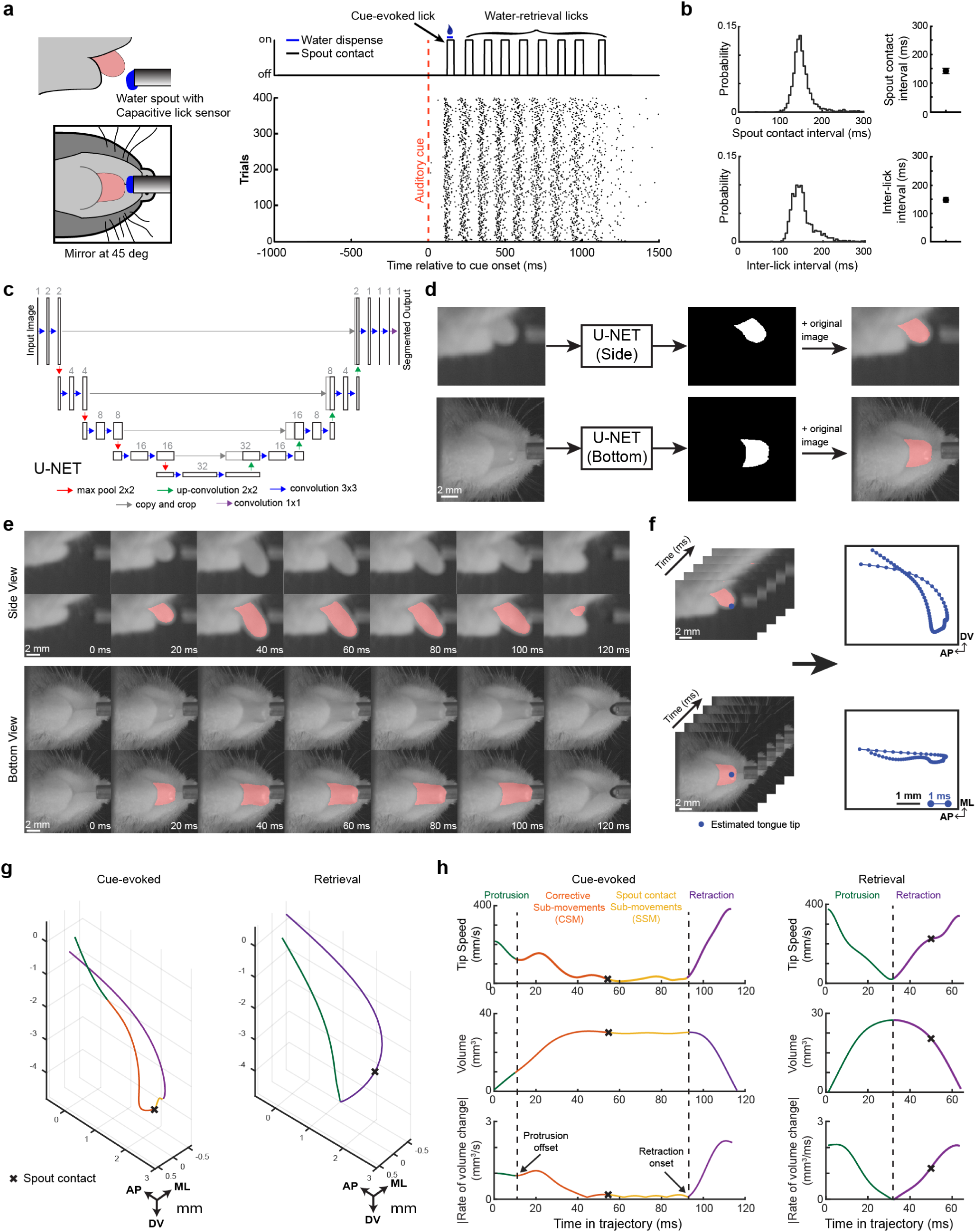
Machine vision based tracking of lingual kinematics identifies corrective submovements important for spout contact. **a**, Left, the tongue was filmed at kilohertz framerate from the side and bottom views during a cued lick task. Right, spout contacts as a function of time for a single trial (top) and lick raster showing spout contact onset times across 400 trials (bottom). **b,** Left: Distributions of inter-spout contact intervals (top) and inter-lick protrusion intervals (bottom) for a single mouse. Right: median±interquartile range (IQR) values across 9 mice. **c,** Architecture of the artificial neural network (U-NET) used to segment the tongue from the background image. U-NET has characteristic symmetrical contraction and expansion paths that simultaneously capture localization and image context. Each box corresponds to a multi-channel feature map and numbers above each layer indicate the number of channels; color-coded arrows indicate sequential processing steps. **d**, Pipeline for tongue segmentation. Left to right, top: side view of the tongue as the input image to U-NET, the identified tongue mask, and the mask plus the input image. Bottom: process is repeated separately for the bottom view of the tongue. **e,** Example frames from side and bottom views across a single lick cycle. Each row shows the raw image above the image overlaid with the U-NET labeled tongue mask. **f,** Tongue tip positions, computed from a 3D tongue model (Extended Data fig. 1), were estimated in each frame (left), resulting in millisecond timescale tracking of tongue tip in two planes. **g,** 3D trajectories of a cue-evoked (left) and water-retrieval lick (right). Protrusion, corrective submovement (CSM), spout submovement (SSM) and retraction phases of the lick are labeled in green, orange, yellow and purple respectively; blue crosses indicate moment of spout contact. **h,** Top to bottom: Tip speed, tongue volume, and absolute value of rate of tongue volume change for the cue-evoked and retrieval licks shown in **g**. Protrusion offsets and retraction onsets were defined as the first and last minima in the rate of volume change (vertical dotted lines). Note that the cue-evoked lick contained CSMs and SSMs between protrusion offset and retraction onset, whereas retrieval licks exhibited a single minimum in rate of volume change, marking the transition from protrusion to retraction.

We defined ‘cue-evoked licks’ as licks initiated before first spout contact and ‘water-retrieval licks’ as licks initiated after the first tongue-spout contact in a bout^16^ (Fig. 1a, g-h). Water-retrieval licks exhibited highly stereotyped kinematics, usually consisting of protrusion immediately followed by a retraction, with no fine-scale submovements in between (Fig. 1g,h, Extended Data fig. 2, Movie S1, Table 1). In contrast, the first cue-evoked lick of each bout (L1) exhibited complex trajectories with longer durations, more acceleration peaks, and more trial-to-trial variability (Fig. 1g,h, Extended Data fig. 2, Table 1). Close examination of cued lick trajectories revealed that the initial tongue protrusion almost never reached the spout (Extended Data fig. 3g, protrusion offset defined as the first minimum in the rate of tongue volume expansion, see Methods). After the protrusion, the animal initiated additional fine-scale tongue submovements prior to retracting. These within-lick submovements, too fast to be seen in real-time videos, were associated with fluctuations in tongue volume and tip speed that were clearly visible in slow-motion (Movie S1). We defined the submovements prior to spout contact as corrective submovements (CSMs) and the submovement after spout contact and before retraction as spout contact submovements (SSMs).

**Fig. 2.**
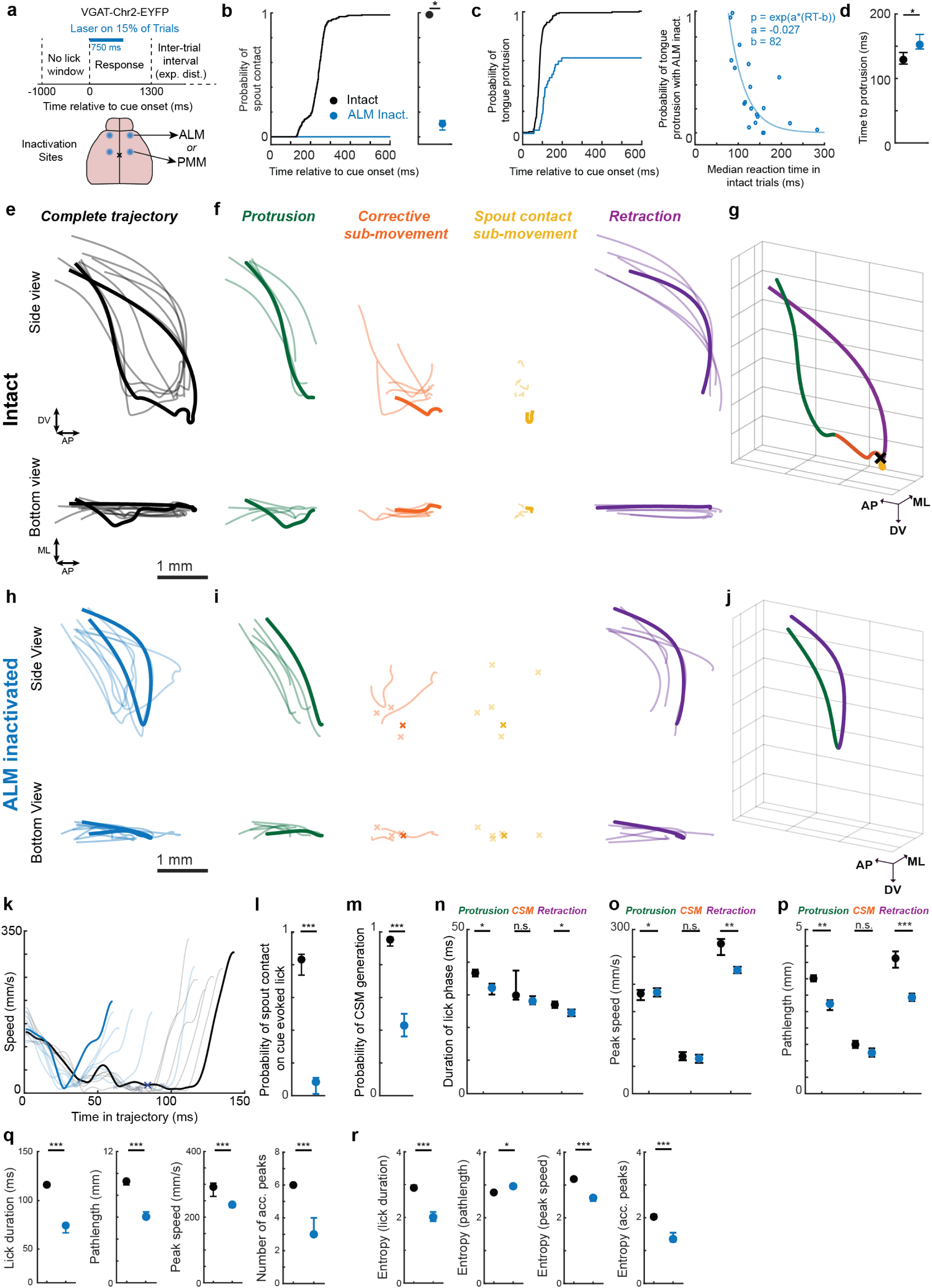
ALM inactivation impairs corrective submovements. **a**, ALM or PMM were bilaterally photoinactivated on 15% of randomly interleaved trials. **b,** Cumulative probability of tongue-spout contact relative to cue onset during control and ALM-inactivated trials (see Extended dat Fig. 8 for PMM results). Right, median±IQR probability of spout contact within a trial across mice (n=20 mice). **c,** Left: data plotted as in **b** for protrusions. Right, protrusion probability during ALM inactivation as a function of reaction time on ALM intact trials; n=20 mice. **d,** Latency from cue onset to tongue protrusion onset in control (black) and ALM-inactivated (blue) trials (median±IQRs, n=20 mice). **e,** Six example tongue tip trajectories from side and bottom views during cue-evoked licks on control trials. A single lick is bold for clarity. **f,** Protrusion, CSM, SSM, and retraction phases of the trajectories from **e** are separately plotted. **g,** 3D trajectory of the bold lick shown in **e-f** with lick phases color-coded. **h-j,** Data plotted as in **e-g** for cue evoked licks on ALM-inactivated trials. x symbols in **i** denote the absence of CSMs and/or SSMs. **k,** Tip speed profiles for the licks from panels **e** and **h**. **l-r,** impact of ALM photoinactivation on L1 spout contact **(l)**, L1 CSMs **(m)**, and the duration **(n)**, peak speed **(o)**, and pathlength **(p)** of distinct lick phases on L1s. **q,** Impact of ALM photoinhbition on the duration, pathlength, speed and number of acceleration peaks in cue evoked licks. **r,** Impact of ALM photoinhbition on the variability of L1 kinematics. Data in **m-r** are median +/-IQRs from trials in which L1 protrusions existed during control (black) and ALM-inactivated (blue) trials with a minimum of 10 data points for each lick phase (*, **, ***, denote p<0.05,0.01 and 0.001 in a Wilcoxon signed rank test; n = 12 mice, all data from sessions with spout at 3.2mm).

**Fig. 3.**
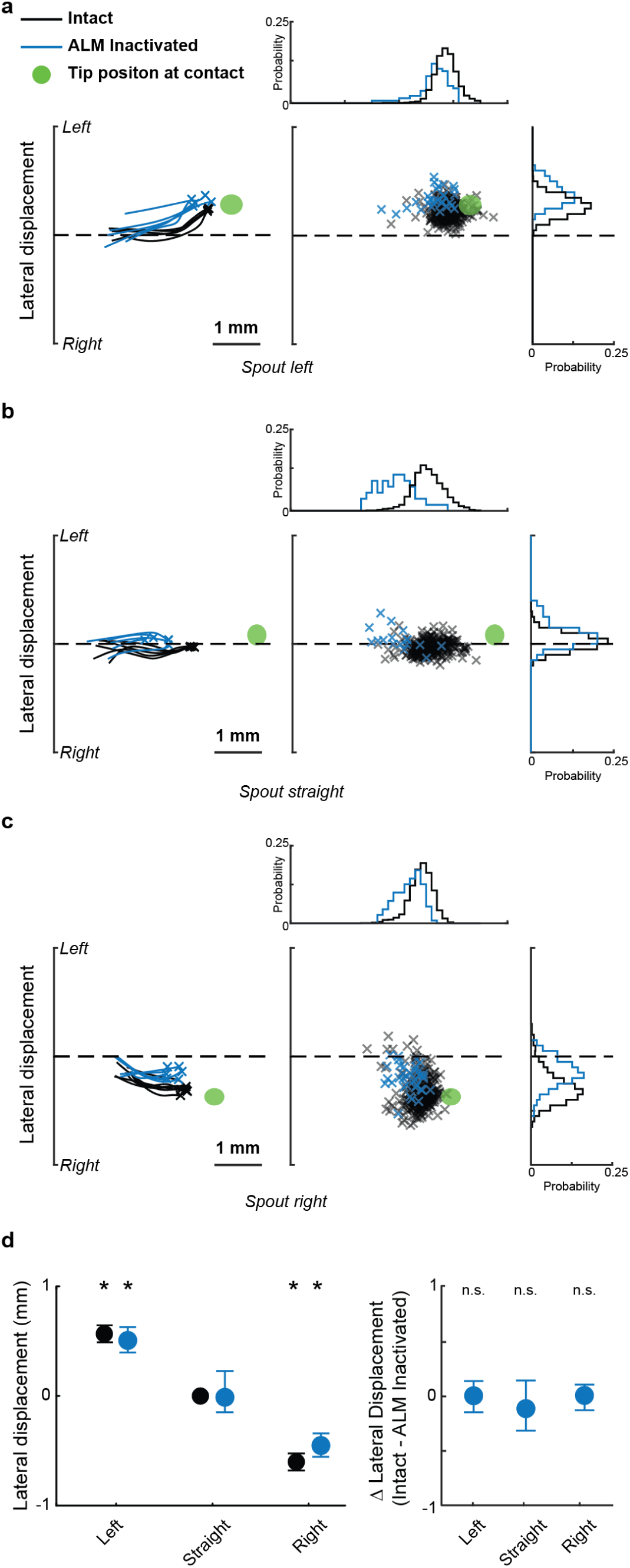
Tongue protrusions remain aimed during ALM inactivation. **a**, Left: Five example tongue protrusions (from bottom view) from a single session with the spout placed to the left (blue, ALM inactivated; black: ALM intact). Green ellipse denotes 95% CI of the tongue tip location at the moment of retraction onset. Center: Scatter plot of tongue tip positions at tongue protrusion offsets. Probability distributions of the ALM intact (black) and inactivated (blue) dots are projected along the axes at top and right (bin size, 120um). Green line, midline. **b,c.** Data plotted in **a** for sessions with centered **(b)** and right **(c)** spout placements. **d,** Left, the lateral placement of the tongue tip at the moment of protrusion offset is plotted across left, straight, and right sessions (black: laser off; blue: ALM Inactivated; data are median±IQR across n=13, 15 and 12 animals for spout left, center and right respectively; * denotes p<0.05 for a Wilcoxon signed rank test). Right, the average difference in lateral displacement between ALM intact and ALM inactivated trials (data are median±IQR displacements across animals n=13,15 and 12 for mice spout left, center and right respectively. n.s. denotes not significant in Wilcoxon signed rank test).

**Table 1.** Kinematics of Cue Evoked and Water Retrieval Licks. Cue-evoked and water retrieval licks had distinct kinematics both at the complete trajectory level and when segmented into lick phases. * denotes a p<0.05, ** denotes a p<0.01 and *** denotes a p<0.001 for a Wilcoxon signed-rank test between cue-evoked and retrieval licks, n = 17 animals.

|  | Cue-Evoked Lick |  | Retrieval Lick |  |
| --- | --- | --- | --- | --- |
|  | Median | IQR [25% 75%] | Median | IQR [25% 75%] |
| *Kinematics of complete lick trajectory* |  |  |  |  |
| Acceleration peaks*** | 6 | [5 6] | 4 | [4 4] |
| Duration*** (s) | 112 | [111 114] | 70 | [68 71] |
| Peak Speed*** (mm/s) | 291.96 | [263.25 303.82] | 314.2732 | [314.17 323.64] |
| Pathlength*** (mm) | 9.28 | [8.71 9.32] | 10.00 | [9.99 10.59] |
| *Entropy of complete lick trajectory* |  |  |  |  |
| Acceleration peaks*** | 2.02 | [2.00 2.10] | 1.28 | [1.27 1.33] |
| Duration*** | 2.98 | [2.90 2.98] | 1.86 | [1.86 1.88] |
| Peak Speed*** | 3.18 | [3.13 3.21] | 2.84 | [2.80 2.84] |
| Pathlength*** | 2.77 | [2.74 2.78] | 2.24 | [2.22 2.40] |
| Probability of generating a CSM*** | 0.95 | [0.91 0.97] | 0.23 | [0.21 0.24] |
| *Kinematics by Lick Phase* |  |  |  |  |
| Duration (ms) |  |  |  |  |
| Protrusion* | 36 | [36 36] | 32 | [30 33] |
| CSM*** | 31 | [30.5 44] | 12.5 | [12.5 13] |
| Retraction** | 27 | [27 27] | 31 | [30 31] |
| Peak speed (mm/s) |  |  |  |  |
| Protrusion*** | 182.76 | [170.97 182.86] | 290.48 | [289.99 294.64] |
| CSM | 67.99 | [64.29 68.92] | 84.39 | [83.56 84.40] |
| Retraction*** | 274.21 | [253.43 276.06] | 315.38 | [310.15 319.11] |
| Pathlength (mm) |  |  |  |  |
| Protrusion** | 3.51 | [3.42 3.52] | 4.32 | [4.59 5.29] |
| CSM** | 1.50 | [1.49 1.57] | 0.68 | [0.62 0.73] |
| Retraction*** | 4.76 | [4.67 4.80] | 5.62 | [5.23 5.73] |

When primates reach to unseen or uncertain targets, CSMs initiated after an initial miss ensure endpoint accuracy, and the number of distinct acceleration peaks in the reach trajectory is correlated with latency to target contact^3, 7^. Similarly, tongue CSMs terminated at precisely clustered tongue tip positions beneath the spout (Extended Data fig. 3) and the number of acceleration peaks per CSM strongly predicted cue-to-spout contact latencies (Extended Data fig. 4a-b). As in primate reach tasks^6^, CSMs depended on target distance: cue-evoked licks to more distant spouts required more CSMs (Extended Data fig. 4c).

**Fig. 4.**
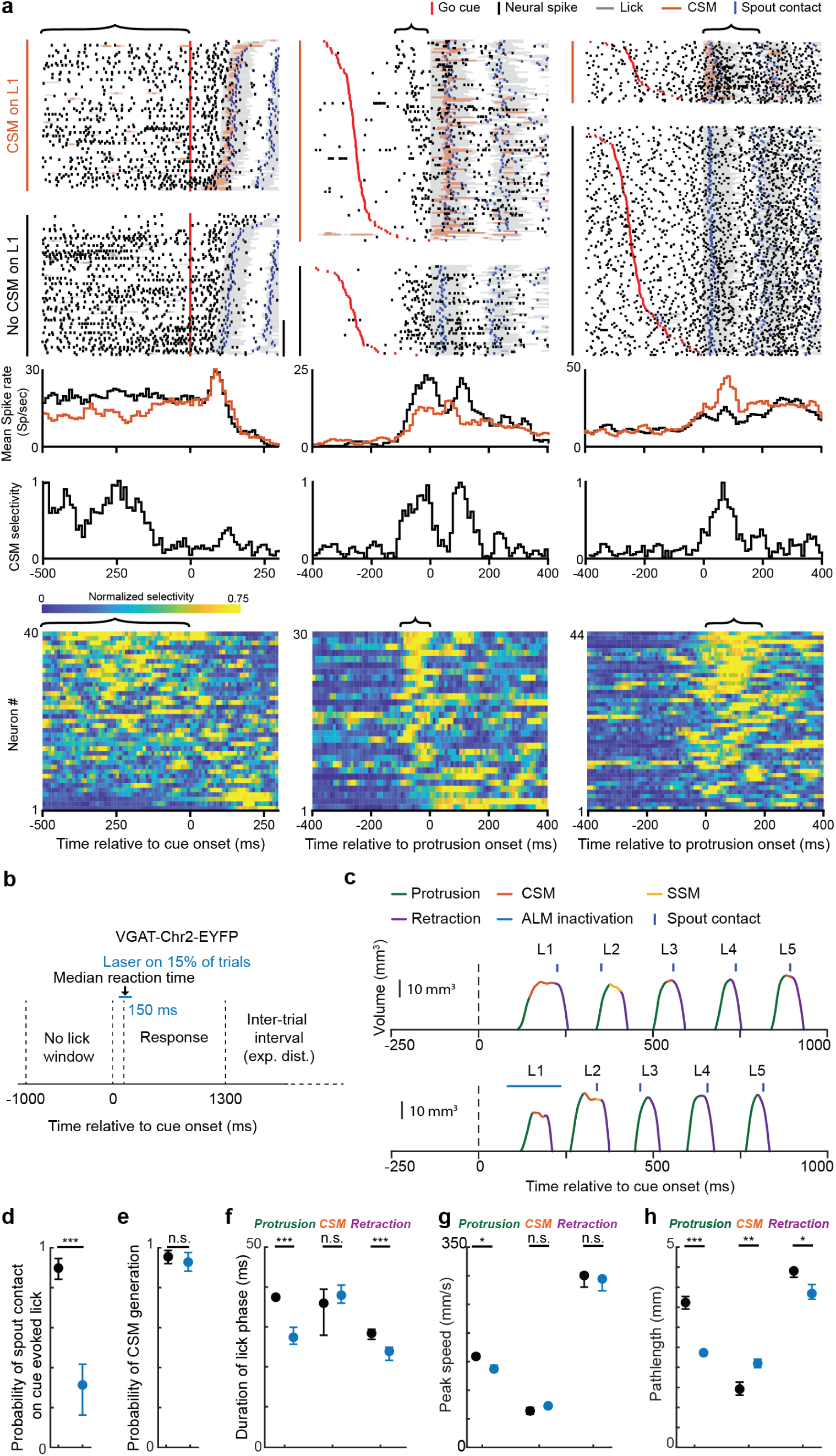
ALM activity reflects upcoming, ongoing and past corrective submovements. **a**, Spike rasters and corresponding rate histograms for three example ALM neurons. Red and blue ticks: go cue and spout contact; Gray and orange shading: Licks and CSMs. Bracket above each raster indicates the window assessed for significant differences in rate histograms between trials with (orange) and without (black) CSMs on L1. Bottom, CSM selectivity, defined as the normalized difference in firing rate from trials with and without CSMs on L1 (Methods). Bottom, ALM population CSM selectivity. Left to right: neurons with CSM selectivity pre-cue, pre-L1, and post-L1. Only neurons with significant trial selectivity are shown (n=40/325 for pre-cue; n=30/325 for pre L1, n=44/325 for post-L1). Note that because each neuron was normalized to its own peak selectivity in the entire trial, the peak selectivity might not lie in the window of interest. **b,** 150 ms duration pulses of photoinhibition were applied 50 ms before the median L1 onset time on randomly interleaved trials. **c,** Tongue volume profiles for a control bout and a bout with pulsed photoinhibition during L1 (blue line at L1 indicates laser-on, resulting in a hypometric L1 that missed the spout. **d-h,** Impact of pulsed photoinhibition during L1 on L1 spout contact **(d)**, L1 CSM generation **(e)**, and duration **(f)**, speed **(g)**, and pathlength **(h)** of L1 lick phases.*, **, ***, denote p<0.05,0.01 and 0.001 in a Wilcoxon signed rank test; n = 12 mice.

To test if CSMs were aimed at spouts or if they were simply random or noisy ‘wiggles’ of the tongue, we studied their kinematics in sessions where spouts were placed at left or right positions (Methods). Both protrusions and CSMs were directionally biased towards remembered spout locations (Fig. 3 and Extended Data fig. 5, Table 2). Together these data show that previously unresolved tongue movements within a lick are controlled and are important for spout contact.

**Fig. 5.**
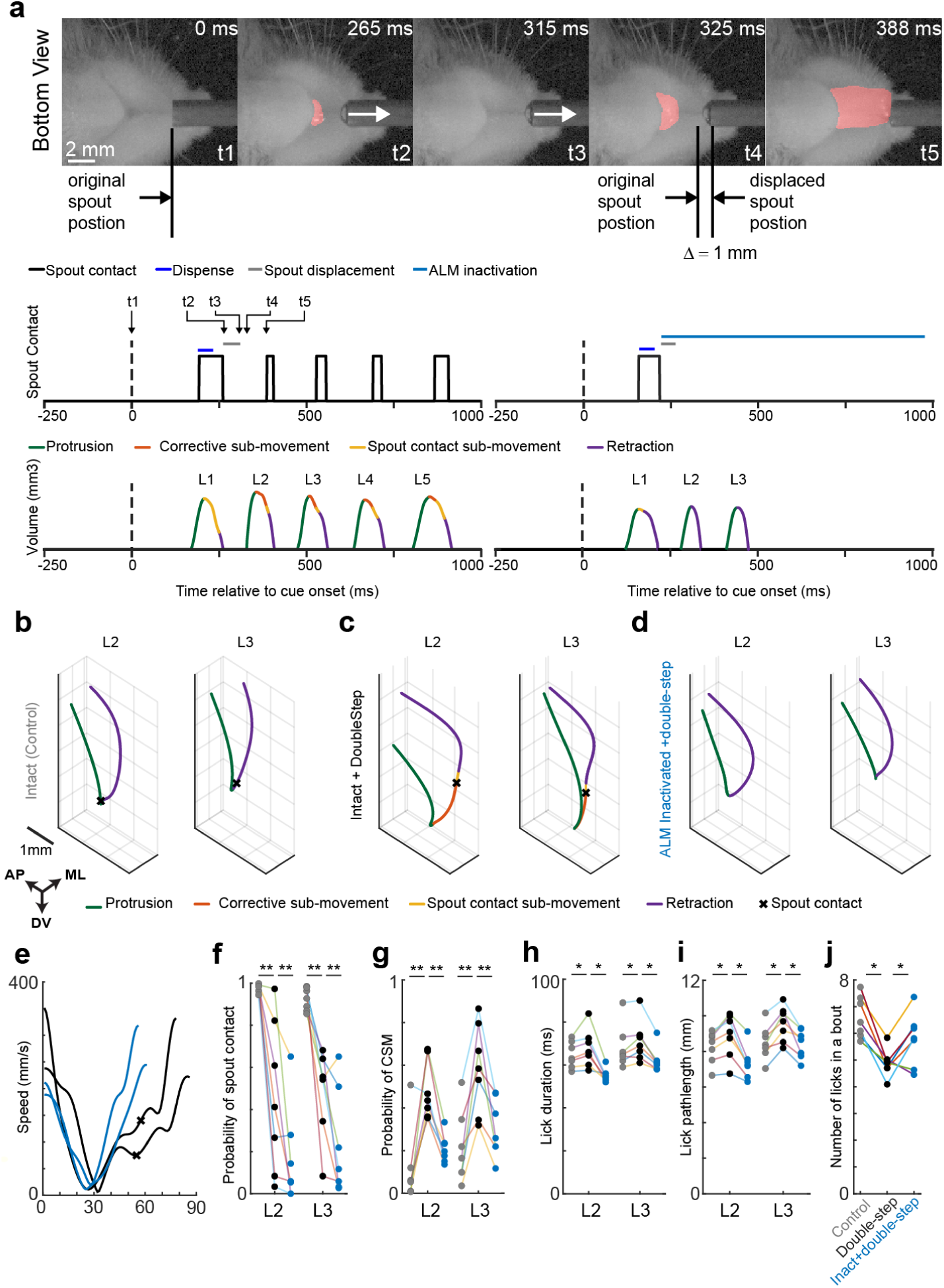
ALM photoinhibition impairs online corrections in double-step experiments. **a**, Top, five still frames from double-step trial showing spout retraction between L1 and L2. Bottom, spout contacts, timesteps from the five still frames, and tongue volumes from a double-step trial with ALM intact (left) and a double-step trial with ALM photoinactivated for 750 ms starting at the moment of L1 contact offset (right, Methods). **b-d,** Tongue tip trajectories during L2 and L3 from control trial (no double-step, no photoinhibition) **(b)**, double-step trial with ALM intact **(c)**, and a double-step trial with ALM inactivated **(d)**. Black x’s denote spout contact. Note ALM photoinhibition impaired CSMs, resulting in hypometric licks that missed the spout. **(e)** Tongue tip speed profiles from licks L2 and L3 in double-step trials from **c-d** (black: control; blue: ALM inactivated; x marks spout contact). **f-g,** Impact of double-step and ALM inactivation on L2 andL3 spout contact **(f)**, CSM probability **(g)**, lick duration **(h)**, lick pathlength **(i)** and number of licks per bout **(j)**. Gray, black and blue circles denote control trials, double-step trials, and double-step+inactivation trials, respectively. *, ** and ** denote p<0.05,<0.01 and <0.001 for a wilcoxon signed rank test; n=7 mice.

**Table 2.** Direction bias of CSM initial velocity vectors. Initial direction of CSM velocity vectors were significantly more biased towards the target in the current session compared to other possible target locations. * denotes p<0.05 for Wilcoxon signed-rank test that the median of direction bias towards the target is significantly different from the median direction bias to the off-target locations. n=13 animals for spout left, n=17 animals for spout straight and n=12 animals for spout right.

| Target | Spout Left |  | Spout Center |  | Spout Right |  |
| --- | --- | --- | --- | --- | --- | --- |
|  | Median | IQR [25% 75%] | Median | IQR [25% 75%] | Median | IQR [25% 75%] |
| Left | 0.79* | [ 0.77 0.84] | 0.47 | [0.47 0.50] | 0.35 | [0.31 0.37] |
| Center | 0.46 | [0.45 0.58] | 0.97* | [0.95 0.97] | 0.60 | [0.42 0.62] |
| Right | 0.42 | [0.40 0.53] | 0.59 | [0.57 0.60] | 0.77* | [0.63 0.79] |

We next wondered why cue-evoked licks contained such prominent CSMs. In primate reaching, uncertainty in the precise location of a target contributes to errors and CSMs^21–23^. If spout position uncertainty contributes to CSMs during licking, then the moment of first tongue-spout contact in a bout could clarify the spout’s position in space and rapidly update the next lick’s motor plan in the same way that, for example, even brief visual feedback at the end of a non-visually guided reach can clarify target position and increase the reach accuracy, reducing the need for CSMs^4, 5, 23^. In some trials, the first one or two licks in a bout failed to make spout contact, providing an opportunity to test how the first contact in a bout affects the next lick. Licks preceding first contact exhibited pronounced CSMs, whereas licks following first contact did not, independent of which lick in a bout made first contact (Extended Data fig. 6a-b, Movie S2). By withholding water until the second spout contact, we additionally observed that both spout and water contact reduced CSM probability in ensuing licks (Extended Data fig. 6c-d). Note that any update to an ensuing lick’s motor plan must rapidly occur in the brief interval between the first spout contact and the ensuing protrusion onset (Latency between spout contact and subsequent protrusion: 94.5ms [87.5 109.5], n = 17 animals). The return to variable cue-evoked licks by the next trial also suggests that uncertainty in the precise location of the spout degrades on second timescales, as observed in memory-guided reach tasks^21–23^. Consistent with this idea, both the probability and duration of CSMs on the first lick in a bout were significantly correlated with the time-since the last spout contact on the previous trial (Extended Data fig. 6c-d, Methods). Together these data suggest that CSMs are prominent when the target location is uncertain, as in primate reaching^21–23^.

**Fig. 6.**
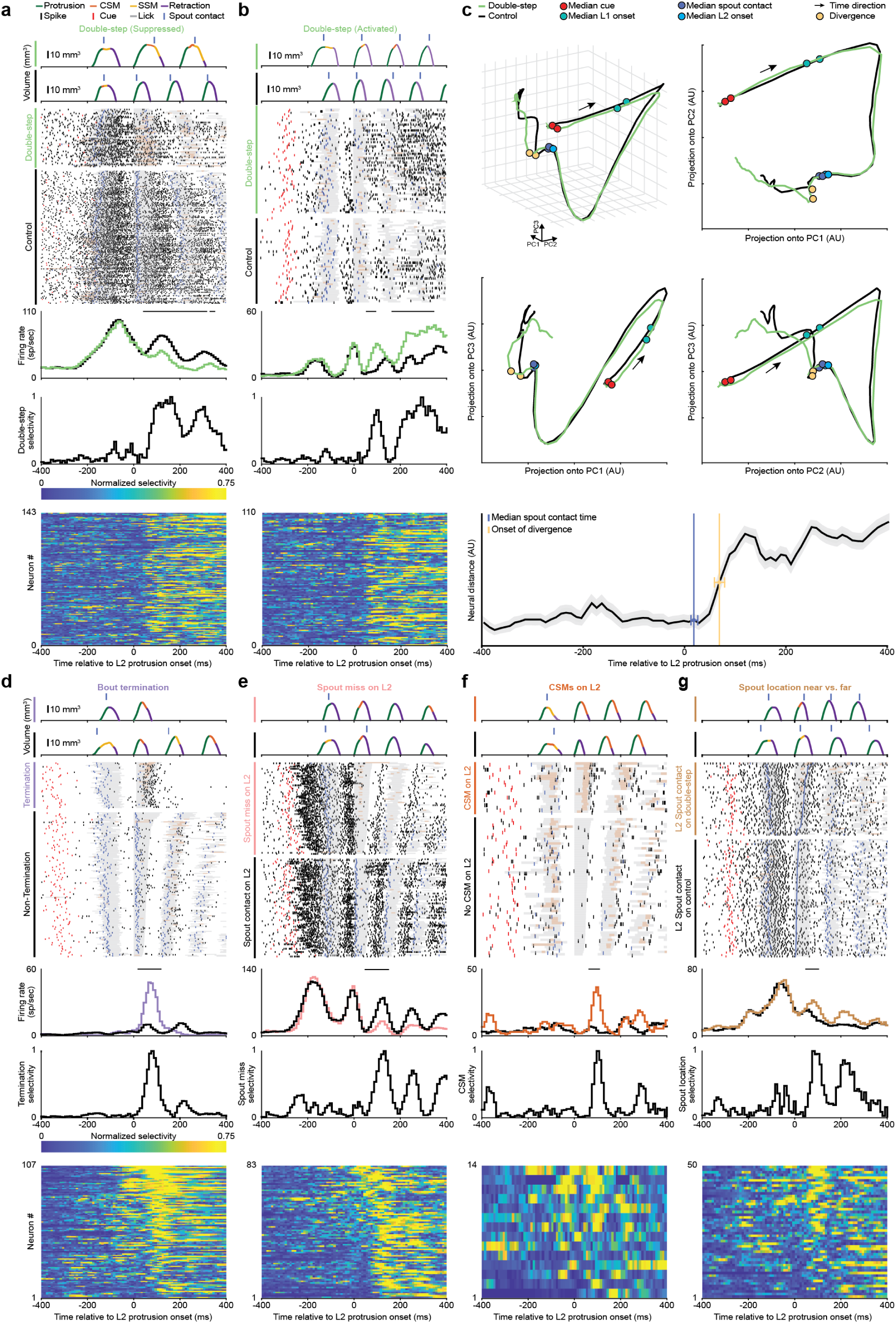
ALM exhibits neural correlates of online corrections in double-step experiments. **a-b**, Tongue volumes, spike rasters, and corresponding rate and double-step-selectivity histograms for 2 example ALM neurons suppressed **(a)** or activated **(b)** by the double-step. Neural activity aligned to L2 protrusion onset. Raster color-codes as in Fig. 4. Bottom, ALM population double-step selectivity, defined as the normalized difference in firing rate from control and double-step trials (Methods). Only neurons with significant trial selectivity are shown (left, n=143/465 activated; right, n=110/465 suppressed on double-step trials). **c,** The first three principal components of ALM population activity during control and double-step trials (black and green, respectively). Bottom, the population difference in firing between control and double-step trials, aligned to L2 protrusion onset. Blue vertical line, median spout contact time on control trials (15 ms [11 21], relative to L2 protrusion onset). Yellow vertical line, time where neural distance reached significance (60 ms [50 70], relative to L2 protrusion onset, Methods). **d-g,** Data plotted as in **(a)** for the following conditions: premature bout termination following L2 (n=107/349 neurons) **(d)**, L2 spout misses resulting in bout continuation (n=83/448 neurons) **(e),** CSMs on L2 misses (n=14/103 neurons) **(f)**, and spout location on L2 contacts (n=50/419 neurons) **(g)**.

### ALM inactivation impairs corrective submovements

To test cortical roles in lingual kinematics, we used VGAT-hChR2-EYFP mice to photoinhibit anterolateral (ALM) or posterior medial (PMM) motor cortical areas, two non-overlapping regions with functional projections to brainstem lingual circuits^11, 24^ (Fig. 2a, Extended Data Fig. 7, and Methods). Photoinhibition was initiated at randomly interleaved cue onsets and lasted 750 ms. Inhibition of ALM, but not PMM, impaired spout contact (Fig. 2b, Extended Data Fig. 8, Table 3-4), consistent with previous studies^12, 15^.

**Table 3.**
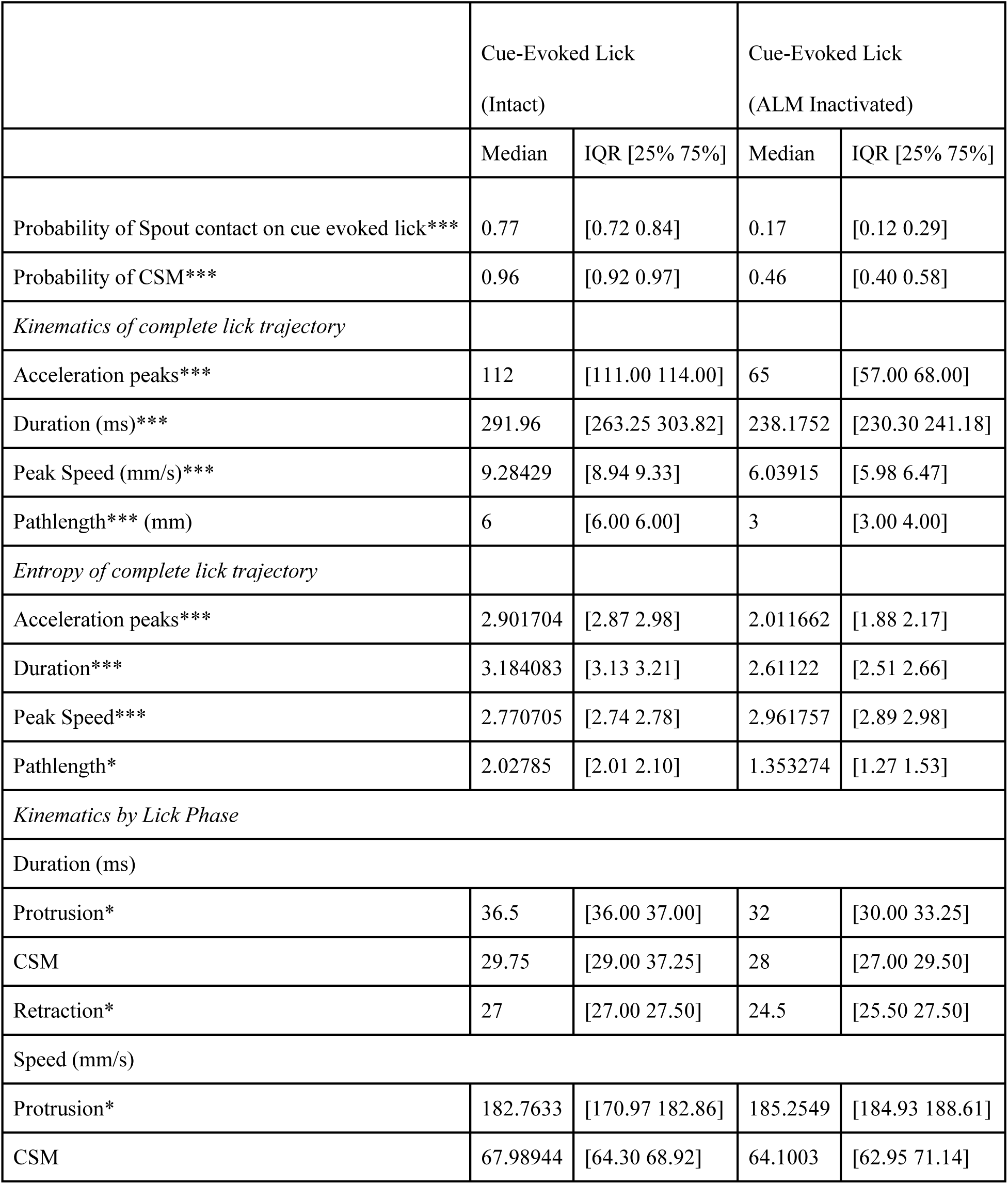
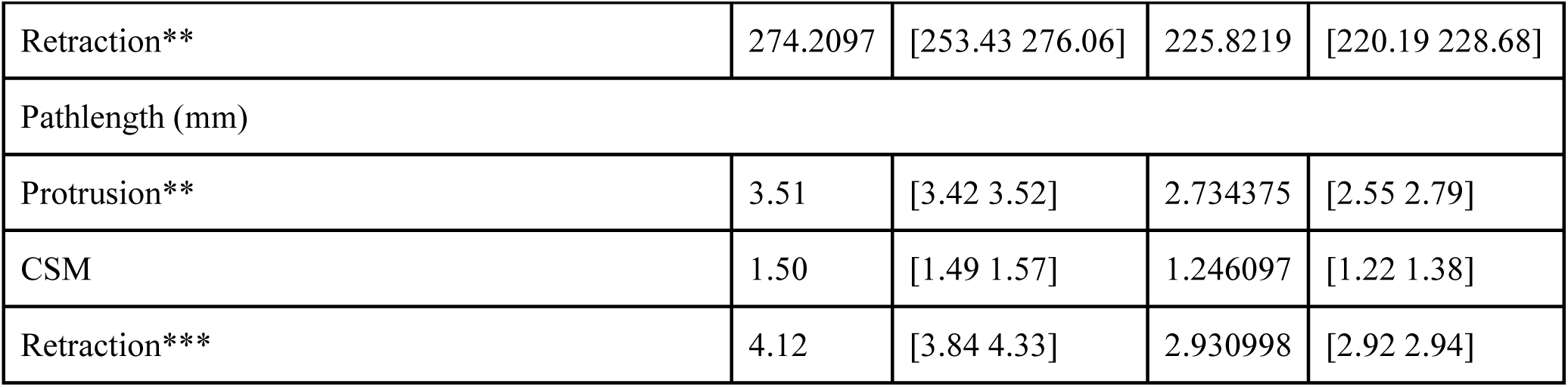
Kinematics of Cue Evoked Licks with ALM inactivation. ALM inactivation altered the kinematics of cue-evoked licks both at the complete trajectory level and when segmented into lick phases. * denotes a p<0.05, ** denotes a p<0.01 and *** denotes a p<0.001 for a Wilcoxon signed-rank test between cue-evoked licks with and without ALM inactivation performed at cue onset, n = 12 animals.

**Table 4.**
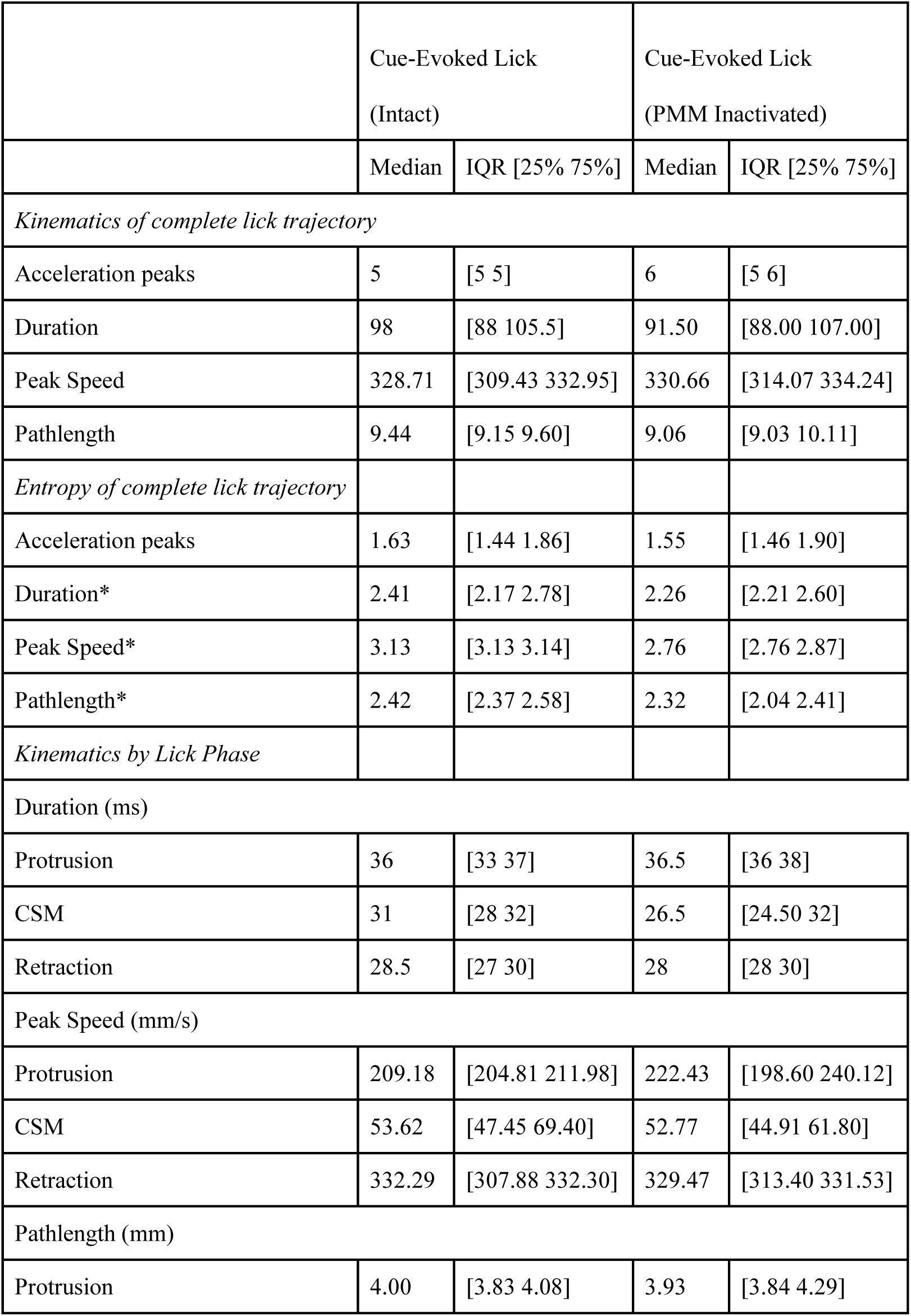
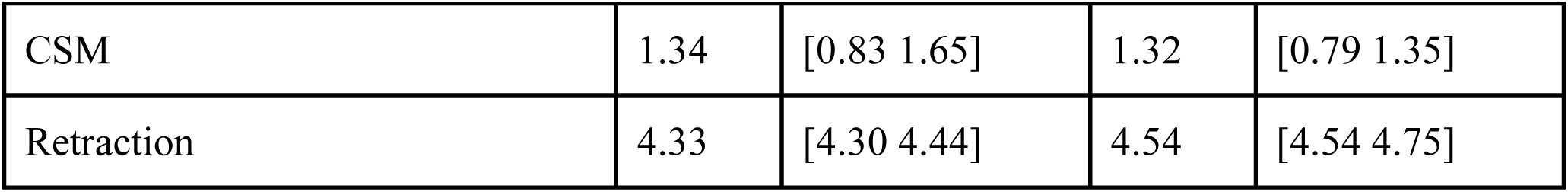
Kinematics of Cue Evoked Licks with PMM inactivation. PMM inactivation did not alter the kinematics of cue-evoked licks both at the complete trajectory level and when segmented into lick phases.* denotes a p<0.05, ** denotes a p<0.01 and *** denotes a p<0.001 for a Wilcoxon signed-rank test between cue-evoked licks with and without PMM inactivation performed at cue onset, n = 9 animals.

We analyzed tongue kinematics on ALM-inactivated trials to examine why spout contact was impaired. ALM inactivation reduced the probability of tongue protrusion in a way that was strongly associated with reaction time (RT). Animals with fast RTs exhibited significantly smaller impairments in protrusion during ALM photoinhibition (Fig. 2c). Cue-evoked licks produced during ALM inactivation were still significantly less likely to make spout contact (Fig. 2b, Movie S3), showing that impaired lick initiation did not fully explain ALM inactivation-associated deficits in spout contact.

Why did cue-evoked licks during ALM inactivation fail to make spout contact even on trials when protrusion occurred? During ALM photoinhibition, cued licks exhibited significantly shorter durations, reduced speeds, reduced pathlengths and fewer acceleration peaks (Fig. 2, Table 3). ALM photoinhibition also reduced trial-by-trial variability of lick kinematics (Methods, Table 3). Critically, on ALM inactivated trials, mice usually failed to produce CSMs and instead immediately retracted the tongue after missed protrusions (Fig. 2m). In rare cases where spout contact occurred during ALM photoinhibition, subsequent water-retrieval licks occurred despite ongoing ALM inactivation (Table 5 and Extended Data fig. 9). Thus, with ALM inactivated, cued licks lacked the CSMs that facilitate spout contact. Within-reach CSMs in primates also rely on cortical activity^25^.

The sparing of tongue protrusions during ALM inactivation led us to hypothesize, first, that protrusions aimed to left or right spouts may not depend on ALM, and second, that ALM inactivations would have a minor impact on performance at very near spout locations, where CSMs are less important for contact. Experiments confirmed these predictions (Fig. 3, Extended Data fig. 10, and Movies S4-S5).

### ALM activity reflects upcoming, ongoing and past corrective submovements

We next recorded ALM activity in sessions with intermediate spout distance (Fig. 4, Extended Data fig. 4c) and compared discharge in trials that lacked or contained CSMs on the first lick of a bout (L1) (325 neurons; 19 sessions; 4 animals). Many ALM neurons exhibited neural correlates of upcoming CSMs before licks were initiated and even before cues (Fig. 4a-b; 40/325 neurons before cue, 30/325 neurons after cue and before protrusion, Methods). Such preparatory activity may reflect the relationship between CSM generation and target uncertainty associated with the intertrial interval (Extended Data fig. 6c-d). ALM activity also reflected ongoing and past CSMs, suggesting additional roles in CSM execution and monitoring (Fig. 4b-c; 44/325 neurons). To test when ALM activity mattered for the initiation and control of CSMs, we briefly photoinhibited ALM for 150 ms starting 50 ms before the median time of L1 protrusion onset (Methods). This intervention, by design, left preparatory activity mostly intact and specifically disrupted activity during L1 (and CSM) execution (Fig. 4d). L1s produced during pulsed inhibition exhibited hypometric protrusions that could be followed by CSMs that usually missed the spout (Fig. 4d-h, Table 6, Movie S6). Together with the inactivations in Fig. 2, these results suggest a role of ALM activity in ongoing licks, but show that protrusions and CSMs can be initiated during ALM inactivation provided preparatory activity is intact.

**Table 5.** Kinematics of Retrieval Licks with ALM inactivation. ALM inactivation made the water-retrieval licks hypometric. * denotes a p<0.05, ** denotes a p<0.01 and *** denotes a p<0.001 for a Wilcoxon signed-rank test between retrieval licks with and without ALM inactivation performed at cue onset. (n=12 animals)

|  | Retrieval Lick<br>(Intact) |  | Retrieval Lick<br>(ALM Inactivated) |  |
| --- | --- | --- | --- | --- |
|  | Median | IQR [25% 75%] | Median | IQR [25% 75%] |
| Probability of spout contact | 0.99 | [0.99 0.99] | 0.971429 | [0.89 0.97] |
| Probability of CSM | 0.26 | [0.15 0.30] | 0.087945 | [0.02 0.13] |
| *Kinematics by Lick Phase* |  |  |  |  |
| Duration (ms) |  |  |  |  |
| Protrusion*** | 32 | [32.00 33.00] | 29 | [28.50 31.00] |
| CSM | 14 | [13.00 16.00] | 13 | [12.00 13.50] |
| Retraction*** | 31 | [30.00 31.00] | 27.5 | [27.50 28.00] |
| Peak Speed (mm/s) |  |  |  |  |
| Protrusion** | 292.32 | [276.68 316.02] | 286.24 | [280.02 287.38] |
| CSM** | 63.55 | [61.45 66.18] | 64.93 | [51.79 64.93] |
| Retraction** | 289.96 | [276.94 302.98] | 248.63 | [247.54 279.64] |
| Pathlength (mm) |  |  |  |  |
| Protrusion** | 4.42 | [4.37 4.65] | 3.52 | [3.44 3.71] |
| CSM | 0.71 | [0.65 0.80] | 0.68 | [0.68 0.78] |
| Retraction* | 5.74 | [5.67 5.98] | 3.86 | [3.81 4.48] |

### ALM photoinhibition impairs online corrections in double-step (target-jump) experiments

In primate reach tasks, CSMs occur in conditions where the likely need for corrections can be planned in advance – such as when a target is uncertain, unseen, or far away^3–7^ – and also in conditions when CSMs are generated on-the-fly, such as in ‘double-step’ experiments when a target unexpectedly jumps mid-reach^8–10, 25, 26^. To disambiguate the roles of cortex in planning corrections in advance versus executing them online, we adapted the ‘double-step’ paradigm to a lick task. Notably, in contrast to primate experiments where animals use visual feedback to detect and correct for target displacement during reaching, our task required corrections to be driven by the absence of a predicted mechanosensory event, the tongue-spout contact. To do this, we detected the offset of tongue-spout contact on L1 in real time and rapidly retracted the spout so that by the onset of the second lick (L2) the spout was at least 1 mm farther away^17, 18^ (Methods). This task tests if mice can implement both within-lick and across-lick corrections. First, to make L2 contact, the tongue might detect a miss and immediately extend substantially farther than usual. To make contact on the third lick (L3), the animal might use the information about L2 outcome to increase L3 pathlength. Finally, following spout misses mice may prematurely terminate the lick bout. With ALM intact, mice exhibited high rates of contact and produced all types of online corrections (Fig. 5, Table 7-8, Movie S7). Both L2s and L3s on double-step trials exhibited increased durations, pathlengths, and CSMs (Fig. 5f-i). Mice also produced less licks per bout (Fig. 5j). Mice can thus produce within- and across-lick adjustments by modifying lick amplitudes to reach farther towards an unexpectedly displaced spout, by producing CSMs, and by prematurely terminating bouts.

**Table 6.** Inactivation at protrusion onset. ALM inactivation at protrusion onset made the cue-evoked licks hypometric. * denotes a p<0.05, ** denotes a p<0.01 and *** denotes a p<0.001 for a Wilcoxon signed-rank test between cue-evoked licks with and without ALM inactivation performed at median protrusion onset. n = 12 animals.

|  | Cue evoked lick<br>(Intact) |  | Cue evoked lick<br>(ALM Inactivation at median protrusion onset) |  |
| --- | --- | --- | --- | --- |
|  | Median | IQR [25% 75%] | Median | IQR [25% 75%] |
| Probability of spout contact *** | 0.95 | [0.92 0.98] | 1.00 | [0.97 1.00] |
| Probability of CSM | 0.90 | [0.84 0.95] | 0.31 | [0.16 0.41] |
| *Kinematics by Lick Phase* |  |  |  |  |
| Duration (ms) |  |  |  |  |
| Protrusion*** | 37.50 | [37.00 38.00] | 27.50 | [26.00 30.00] |
| CSM | 36.00 | [28.00 42.00] | 38.00 | [36.50 40.50] |
| Retraction*** | 28.50 | [27.00 29.50] | 24.00 | [21.75 25.00] |
| Peak Speed (mm/s) |  |  |  |  |
| Protrusion* | 159.02 | [153.29 163.06] | 137.04 | [133.91 146.91] |
| CSM | 63.97 | [58.23 69.72] | 72.75 | [71.71 75.50] |
| Retraction | 300.40 | [280.23 305.23] | 294.58 | [273.66 297.95] |
| Pathlength (mm) |  |  |  |  |
| Protrusion*** | 3.62 | [3.45 3.77] | 2.36 | [2.30 2.37] |
| CSM** | 1.46 | [1.31 1.63] | 2.10 | [1.99 2.20] |
| Retraction* | 4.41 | [4.25 4.46] | 3.84 | [3.70 4.07] |

**Table 7.** Kinematics of L2 licks during double-step and double-step with ALM inactivation. L2 licks exhibited CSMs on double step trials that were dependent on ALM activity. * denotes a p<0.05, ** denotes a p<0.01 and *** denotes a p<0.001 for a Wilcoxon signed-rank test between L2 licks in control and control with double step. ^+^ denotes a p<0.05, ^++^ denotes a p<0.01 and ^+++^ denotes a p<0.001 for a Wilcoxon signed-rank test between L2 licks control with double step and doublestep with ALM inactivation. (n=7 animals)

|  | L2<br>(Intact, no double step) |  | L2<br>(Intact, double step) |  | L2<br>(ALM inactivated, double step) |  |
| --- | --- | --- | --- | --- | --- | --- |
|  | Median | IQR [25% 75%] | Median | IQR [25% 75%] | Median | IQR [25% 75%] |
| Probability of spout contact** ++ | 0.99 | [0.97 0.99] | 0.41 | [0.13 0.77] | 0.06 | [0.01 0.24] |
| Probability of CSM** ++ | 0.05 | [0.02 0.11] | 0.43 | [0.37 0.62] | 0.20 | [0.16 0.23] |
| *Kinematics of complete lick* |  |  |  |  |  |  |
| Duration (ms)* + | 63.00 | [59.75 70.25] | 66.00 | [61.13 72.00] | 54.50 | [53.25 55.75] |
| Pathlength (mm/ms)* + | 8.56 | [7.66 8.89] | 9.09 | [8.05 9.92] | 7.23 | [6.71 7.52] |
| *Kinematics by Lick Phase* |  |  |  |  |  |  |
| Duration (ms) |  |  |  |  |  |  |
| Protrusion | 31 | [30.00 32.00] | 31 | [31.00 31.50] | 28 | [28.00 28.00] |
| CSM | 13 | [10.00 13.50] | 15.5 | [14.50 16.50] | 12 | [11.50 13.50] |
| Retraction<sup>+</sup> | 28 | [26.00 28.00] | 26.5 | [26.50 30.00] | 25 | [24.00 25.00] |
| Peak Speed (mm/s) |  |  |  |  |  |  |
| Protrusion<sup>+</sup> | 261.57 | [249.27 274.29] | 249.65 | [243.10 268.95] | 248.79 | [242.02 248.81] |
| CSM | 70.27 | [49.38 75.21] | 76.75 | [75.55 78.65] | 78.72 | [70.94 80.02] |
| Retraction*<sup>+</sup> | 271.58 | [269.86 274.46] | 287.81 | [287.70 291.42] | 269.58 | [269.58 279.30] |
| Pathlength (mm) |  |  |  |  |  |  |
| Protrusion<sup>+</sup> | 3.70 | [3.63 3.93] | 3.69 | [3.48 3.85] | 3.12 | [3.07 3.19] |
| CSM | 0.57 | [0.51 0.63] | 0.66 | [0.60 0.72] | 0.60 | [0.55 0.60] |
| Retraction<sup>+</sup> | 4.46 | [4.37 4.55] | 4.55 | [4.21 4.62] | 3.96 | [3.95 4.00] |

**Table 8.**
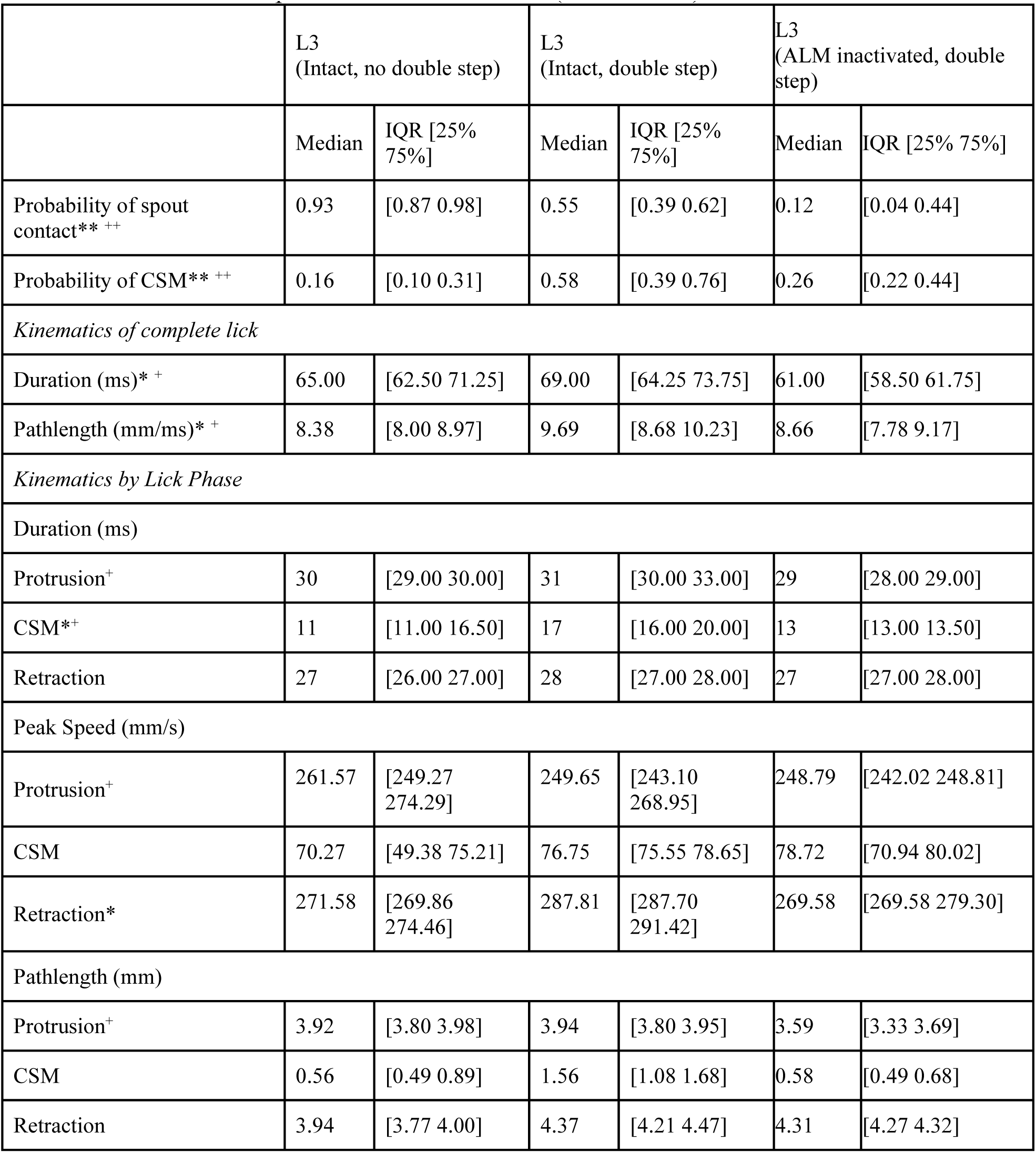
Kinematics of L3 licks during double-step and double-step with ALM inactivation. L3 licks exhibited CSMs on double step trials that were dependent on ALM activity. * denotes a p<0.05, ** denotes a p<0.01 and *** denotes a p<0.001 for a Wilcoxon signed-rank test between L3 licks in control and control with double step. ^+^ denotes a p<0.05, ^++^ denotes a p<0.01 and ^+++^ denotes a p<0.001 for a Wilcoxon signed-rank test between L3 licks control with double step and doublestep with ALM inactivation. (n=7 animals)

To test the role of ALM in these corrections, we photoinhibited ALM for 750ms right after L1 withdrew from the spout on a randomly interleaved subset of double-step trials (Methods). ALM inactivation impaired L2 and L3 contact and significantly reduced L2 and L3 durations, pathlengths, and CSMs (Fig. 5e-i. Tables 7-8, Movie S8). ALM photoinhibition also significantly prolonged lick bouts - as if the mouse did not detect spout misses (Fig. 5j, Tables 7-8). ALM activity is thus important for on-the-fly corrections produced both within and across licks in a bout.

### ALM exhibits neural correlates of online corrections in double-step experiments

Several neural signals are important for these corrections. First, making a correction on L2 requires signals associated with the absence of a predicted spout contact and with the motor correction. Updating a plan for L3 after an L2 miss also requires signals associated with the absence of a predicted spout contact, whereas updating a plan for L3 after L2 contact requires information about whether or not the spout was near (on control trials) or far (on double-step trials). Such spout position-dependent discharge would require ALM to integrate the mechanosensory event of contact with proprioceptive information about tongue posture at the moment of contact. To our knowledge it remains unknown if the motor cortex in any species exhibits such position-at-contact signals unrelated to reward or visual feedback.

We recorded ALM activity in double-step sessions (n=465 neurons, n=28 sessions, n= 4 animals) and discovered that many neurons discharged differently on control versus double-step trials (253/465 neurons). Principal component analysis revealed that the population activity on double-step trials diverged from control trials in the middle of L2 (time of activity divergence: 60+/-10 ms after L2 protrusion onset; duration of L2 on double-step trials: 66 ms [57 81]), fast enough to play a role in online corrections^8–10^. Closer examination of single neuronal discharge on double-step trials revealed neural correlates of premature bout termination following spout misses^18^ (Fig. 6d, 107/349 neurons), spout misses even when lick bouts continued (Fig. 6e, 83/448 neurons), and CSMs during L2 (Fig. 6f, 14/103 neurons). Many neurons also encoded spout location: tongue-spout contact at far locations resulted in significantly different discharge than tongue-spout contact at near locations, even though both contacts were identically rewarded (Fig. 6g, 50/419 neurons). Similar error- and correction-related activities were observed on L3 (Extended Data Fig. 11). These data show that ALM exhibits signals associated with both errors and corrections with appropriate timing to influence ongoing licks.

High speed videography revealed novel aspects of tongue control in bats, hummingbirds, chameleons, cats and bees^27–30^. Here we discovered that single licks exhibit complex trajectories with hallmarks of online motor control, including motor cortex-dependent corrective submovements that facilitate target contact. Interestingly, as in primate reach tasks, CSMs predicted target contact time, depended on target distance and occurred when target location was uncertain or unexpectedly displaced^3–7^. Also as in primates, motor cortical activity was important for successful correction and reflected upcoming, ongoing, and past corrections^8–10, 25, 26^. We also discover that motor cortex can signal position-at-target-contact signals consistent with integration of mechanosensory and proprioceptive inputs, and predict such signals will exist in motor cortical areas of other species in tasks that require contact-driven corrections.

Comparative approaches distinguish general principles from behavior-, effector-, and species-specific solutions to motor control problems. Our discovery that mouse tongue, a muscular hydrostat with no joints, and primate limb adhere to common control principles suggest canonical roles of cortex in error corrections important for the accuracy of ongoing movements, including lingual trajectories important for coherent speech^2^.

## Methods

### Animals and Surgery

All experiments were carried out in accordance with NIH guidelines and were approved by the Cornell Institutional Animal Care and Use Committee. Twenty six VGAT-ChR2-EYFP (Jackson Laboratory, JAX Stock #014548) and four C57/B6J (Jackson laboratory, JAX Stock #000664) animals > 16 weeks of age were individually housed under a 12hr light/dark cycle for the duration of the study, and were tested during the dark phase. On days when mice weren’t being trained or tested, mice received 1 ml of water. Mice were trained and tested in experimental sessions that lasted 0.5h to 1h. If the mice did not receive at least 1ml of water during the behavioral session, their water was supplemented to meet the 1ml/day requirement.

Mice were deeply anesthetized with isoflurane (5%). Fur was trimmed, and mice were placed in a stereotaxic frame (Kopf Instruments, Tujunga, CA). A heating pad was used to prevent hypothermia. Isoflurane was delivered at 1-3% throughout surgery; this level was adjusted to maintain a constant surgical plane. Ophthalmic ointment was used to protect the eyes. Buprenorphine (0.05 mg/kg, subcutaneous) was given before the start of surgery. A mixture of 0.5% lidocaine and 0.25% bupivacaine (100 μL) was injected subdermally along the incision line. The scalp was disinfected with betadine and alcohol. The scalp was then removed with surgical scissors to expose the skull, which was thoroughly cleaned.

For optogenetic experiments, four craniotomies were made over ALM (2.5 A/P±1.5 M/L) and PMM (0.5 A/P±1.5 M/L). A 400µm optical fiber embedded in a 1.25 mm metal ferrule (Thor Labs Inc., Newton, NJ) was then implanted bilaterally above these areas and held in place with a layer of Metabond (Parkell, Inc., Edgewood, NY). Mice were then implanted with a custom modified RIVETS headplate for head restraint during the behavioral sessions. Another layer of Metabond was applied to firmly hold the implants in place, and the surrounding skin was sutured.

For acute electrophysiology experiments, a craniotomy was made over visual cortex (-3.5 A/P±3 M/L), along with two fiducials that were made bilaterally over ALM and marked with black ink. A ground screw (W.W. Grainger, Inc., Lake Forest, IL) soldered to a gold pin (A-M Systems, Sequim, WA) was then screwed into the craniotomy and a headplate was secured to the skull. The skull was then covered with a thin layer of clear Metabond. Post-operative enrofloxacin (5 mg/kg), carprofen (5 mg/kg), and lactated ringers (500 μL) were administered subcutaneously.

### Behavior

To simultaneously image two orthogonal planes of the mouse’s orofacial movements, we placed a mirror (Thorlabs ME1S-P01 1”) angled at 45 degrees below the mouse’s mouth. We used a Phantom VEO 410L camera with a Nikon 105mm f/2.8D AF Micro-Nikkor lens to acquire videos with a resolution of 192x400 pixels at 1000 fps. Custom LabVIEW code for behavioral training was run on a training system built using a National Instruments sbRIO-9636 FPGA. Details regarding the behavioral rig, including parts list, diagrams, and instructions, can be found on our Github page (https://github.com/GoldbergLab). Briefly, behavior rigs consisted of custom 3D-printed clamps that were used for head-fixation, an audio system for generating cues (Med Associates) and a blue LED that served as a masking light for optogenetics. A 0.0072-inch stainless steel lickport was used to record spout contacts using a capacitive sensor (Atmel) and deliver water rewards via a solenoid valve (The Lee Company). We used a custom circuit (Janelia Farms, HHMI) to detect spout contact onsets in place of the capacitive lick sensors during electrophysiology experiments to reduce noise. For double-step experiments, lickports were attached to a servomotor (Faulhaber) which was used to retract the spout by a pre-calibrated distance (1 mm or 4 mm) at L1 spout contact offset.

### Behavioral Training

Five days after surgery and postoperative recovery, mice were started on water restriction. Mice were restricted to 1mL of water per day and their body weight was recorded daily. The behavioral training began after mice reached a steady state of body mass of 80% their original body weight with water restriction. Mice typically reached the steady state body weight in 5-6 days. In the first behavioral sessions, mice were head restrained and water (3uL per dispense) was delivered paired with an auditory cue (3.5 kHz). The spout was placed directly ahead of the mouse, approximately 1.6mm from the mouse’s incisors. The auditory cues had an inter-trial interval with an exponential distribution, which provided a flat hazard rate such that the probability of a cue was not altered over the duration of the trial. After the mice learned to reliably lick the spout following the auditory cue, we imposed a 1 second no spout contact window prior to the onset of the auditory cue. If the animal made spout contact within this window, the inter-trial interval was extended by an interval randomly drawn from the exponential distribution. This discouraged mice from spontaneously licking the spout and ensured that the licking we observed was in response to the auditory cue. Water delivery in subsequent sessions was made contingent on spout contact within 1.3 seconds of the auditory cue. Mice were considered to have reached criterion once performance reached >95% in the task and the proportion of trials with premature licking was less than ∼10%, with little if any licking during the inter-trial interval. Once trained with the spout at 1.6mm, photoinhibition experiments were completed if required (Extended Data Fig. 10), and the spout was moved back to 3.2mm from the mouse’s incisors. Mice were trained with the spout at 3.2 mm for 1 - 2 sessions, then inactivation experiments were performed either at cue-onset or 50ms before the median time of L1 protrusion onset calculated from the previous session. The spout was then placed approximately 60 degrees from midline to the left or right of the mouse (counter-balanced) at a distance of 3.2 mm. Mice were trained in this first direction for several weeks (typically 14 days) before inactivation experiments were performed. This procedure was repeated for the remaining direction.

### Photoinhibition

We used laser LED light sources (LDFLS_450-450, Doric Life Sciences Ltd), attached to an optical rotary joint (FRJ_1×2i_FC-2FC_0.22) and delivered light to the implanted cannulas using 400 µm, 0.43NA lightly armored metal jacket patch cords. The light sources were set to analog input mode and driven with a square (10mW) or sinusoidal pulse (40 Hz, 10mW peak). For inactivations performed at cue-onset, the duration of inactivation was 750 ms with a 100 ms ramp-down^31^. For inactivations performed at L1 protrusion onset, we imaged the tongue as mice were performing the task one day prior to inactivation. We then calculated the median L1 protrusion onset time individually for all mice, and inactivated ALM for 150 ms with a 100 ms linear ramp-down starting at 50ms before the median L1 protrusion time. For double-step experiments, inactivations (750 ms duration with a 100 ms ramp down) were initiated at the moment of L1 spout contact offset detected in real time.

### Electrophysiology

Extracellular recordings were made acutely using 64-channel silicon probes (ASSY-77 H2, Cambridge Neurotech). 64-channel voltage signals were amplified, filtered and digitized (16 bit) on a headstage (Intan Technologies), recorded on a 512-channel Intan RHD2000 recording controller (sampled at 20 kHz), and stored for offline analysis. 12-24 hours prior to recording, a small (1.5 mm diameter) craniotomy was made unilaterally over ALM centered on the fiducial. The probes were targeted stereotaxically to ALM, lowered to a depth of 1000 - 1100 microns. Recording depth from the pial surface was inferred from micromanipulator readings. To minimize brain movement, 1.8% low-melt agarose (A9793-50G, Sigma Aldrich) in 1x phosphate-buffered saline (Corning) was pipetted in the craniotomy following probe insertion. Three to seven recordings were made from each craniotomy. After each recording session, the craniotomy was filled with silicone gel (Kwik-Cast, World Precision Instruments). Carprofen (0.05 mg/kg) was given daily to reduce inflammation.

### Artificial Deep Neural Network for Segmentation

We used an implementation of a semantic segmentation neural network (U-NET) to identify and segment the tongues from high speed videography. U-NET uses a contracting path that is thought to identify context, i.e. “is the tongue present in this image?”, and a symmetric expanding path that precisely localizes the relevant object, i.e. “where is the tongue present in the image?”

#### U-NET Architecture

The contracting path of the network was constructed as a series of 5 repeating modules. Each module was an application of two 3x3 convolutions, with each convolution followed by a ReLU and 2x2 max pooling operation with stride 2 for downsampling. At each downsampling, the number of feature channels was doubled. The number of channels for the first module was 2, and thus for the remaining modules they were 4, 8, 16 and 32 respectively. Dropout of 0.7 was added at the output of module 4 and 5. The expanding path of the network was symmetric to the contracting path, with 4 repeating modules. Each module had: First, a 3x3 convolution with half the number of channels from the previous module. Second, an upsampling step that doubled the frame size. Third, a concatenation step that merged the output of the current module with that of the symmetric module from the contracting path. And finally, two 3x3 convolutions, with each convolution followed by a ReLU. The last layer of the network was a 1x1 convolution layer that followed the last layer of the expanding path. This network had a sigmoid activation function and gave the probability of an individual pixel being a part of the tongue.

#### Network training

The network was trained on 3668 frames pseudo randomly selected from the dataset of 25,258,017 frames from 12 animals across all their sessions. The training set was balanced such that half of the 3668 frames contained visible portions of the tongue. The frames were then manually annotated with both the side view and the bottom view using a custom GUI. Separate networks were trained for the side and bottom views. The networks were trained with a batch size of 256 images, using the ‘adam’ optimizer and a binary cross entropy loss function. The networks were trained till the loss function reached an asymptotic value of 0.0047 for the side view network and 0.0023 for the bottom view network, with a validation accuracy of 0.9979 and 0.9991 respectively. Both networks reached asymptotic performance within 4000 epochs. In order to find the ideal architecture, we performed hyperparameter optimization with the scale of the network and the dropout rate as the two axes. We found that there was no statistical difference in the binary cross entropy loss between the largest (first bank = 256 layers) and the smallest (first bank = 2 layers) networks we tested. There was also no statistical difference in the loss for the dropout rates we adopted. For our purpose, we chose the networks with the least loss that consistently converged.

### Extracting 3D tongue kinematics

To obtain the full 3D kinematics of the tongue tip during a lick bout, we performed a visual hull reconstruction using two orthogonal views (bottom and right side) of the tongue filmed via high-speed videography. This hull reconstruction procedure is contingent upon crisp 2D silhouettes of the tongue from both the bottom and side views, which were obtained by U-NET segmentation. We next constructed a 3D voxel representation of the tongue by identifying voxels that map onto the tongue silhouette when projected back into the 2D image space. Intuitively, this can be thought of as placing the bottom and side view images on adjacent faces of a cube, projecting the silhouettes in towards the center of the cube, and identifying the 3D intersection of these projections (Extended Data fig. 1a). For trials in which the side view of the tongue tip is occluded by the lick spout, we estimate the shape of the occluded tongue region by fitting a cubic spline to the boundary of the side silhouette and extrapolating the boundary spline into the occluded region.

We obtained 3D coordinates of the tongue centroid by averaging, and then defined the tongue tip as the position on the tongue that is furthest from the centroid in the direction of the lick, which we located using a two-step search process (Extended Data fig. 1b). In the first step, we defined an initial search vector, which points forward (anterior) and down (ventral) from the tongue centroid. This initial search vector was used across all videos. Using this initial search vector, we identified voxels in the tongue hull that satisfied the search criteria of i) the vector connecting the voxel to the centroid made an angle of less than 45 degrees with the initial search vector and ii) the distance from the centroid to the voxel was ≥75% of all voxel-to-centroid distances. We took the collection of voxels that satisfy these criteria, which we called candidate voxels, and calculated their mean location. The unit vector between the tongue centroid and the mean location of the candidate voxels was then used as the search vector for the second step of the search process, as it pointed in the rough direction of tongue tip. The second step of the search process followed a similar pattern to refine the search described above. Using the refined search vector from step one, we performed a search for voxels that were i) within a given angular range (15 degrees) of the search vector and ii) were located on the boundary of the tongue hull. The average location of this second set of candidate voxels was defined to be the tongue tip (Extended Data fig 1c). The resultant 3D kinematics for the tongue tip were filtered using an 8 pole, 50 Hz low-pass filter.

### Trajectory Analysis

Tongue volume was determined from the convex hull reconstruction from the segmented images (see “Extracting 3D tongue kinematics”). Tongue tip trajectories were segmented into three distinct phases on the basis of the rate of volume change of the tongue. The protrusion phase was defined as the time from when the tongue is detected up to the first minimum in the rate of volume expansion of the tongue. The retraction phase was defined as the time from the last minima of the rate of volume expansion of the tongue until the tongue was back in the mouse’s mouth. We further defined the movements prior to spout contact and after protrusion as corrective submovements (CSMs) and the submovement after spout contact and before retraction as spout contact submovements (SSMs).

Instantaneous speed was calculated as a one-sample difference of the position vector and pathlength was calculated as the cumulative sum of the one-sample difference of the position vector over the entire trajectory. Acceleration was calculated as the one sample-difference of the instantaneous speed. Peaks were identified using the findpeaks function in MATLAB. Lateral displacement was defined as the distance of the tip position from the midline of the mouse. The midline of the mouse was defined as the line that passes through the point equidistant between the mice’s nostrils and the midpoint of the mouse’s incisors. Entropy for the kinematic parameters was calculated as -ΣP_i_*log(P_i_) where P was the probability of the kinematic parameter being in bin **i**. We used bin sizes of 5ms, 100um and 5mm/s for durations, path lengths and peak speeds respectively.

Direction bias was estimated as the dot product of the initial CSM direction vector and either the target direction vector or the simulated off-target direction vector. The CSM direction vector was defined as the direction vector from the location of the tongue tip at the onset of the CSM to the location of the tongue tip at the first speed minimum. The target direction vector was defined as the direction vector from the location of the tongue tip at the onset of the CSM to the median location of the tongue tip at retraction onset in that session. Similarly, the simulated off-target direction vector was defined as the direction vector from the location of the tongue tip at the onset of the CSM to the simulated target locations in that session.

Since targets were changed across sessions and not within sessions, and not all mice were trained in all directions, the simulated off-target direction vectors were defined as follows. For the left sessions: the center/straight simulated target had the same A/P location and was on the midline. The right simulated target had the same A/P location as the target, and symmetrical M/L location from the midline. For example, if the left target was at +4mm A/P and +1.2mm M/L, the right simulated target would be at +4mm A/P -1.2mm M/L and the center straight target would be at +4mm A/P 0 M/L. For the right sessions: symmetrical to the left sessions. For the center sessions: Both the right and left simulated targets had the same A/P location, and M/L was ±1.2mm.

### Electrophysiology Analysis

Extracellular voltage traces were first notch-filtered at 50/60 Hz. The data were then spike sorted automatically with Kilosort (https://github.com/cortex-lab/Kilosort), and curated manually with Phy2 (https://github.com/cortex-lab/phy). During manual curation, units containing low-amplitude spikes and/or non-physiological or inconsistent waveform shape were discarded and not included in further analyses. Neurons with less than 10 trials in any of the conditions tested were excluded for all analyses performed below.

To determine if a neuron’s firing rate was significantly correlated to L1 CSMs (Fig. 4), we assessed the difference in firing rate between L1 CSM-containing and L1 CSM-lacking trials in three epochs: Before the cue onset, before the onset of the lick protrusion and after the onset of lick protrusion. To identify pre-cue L1 CSM responsive neurons, we aligned the recordings to cue onset and assessed significance in spike counts across the trial types in the time period from 500 ms before cue onset to cue onset (WRS test, p<0.05). To identify peri-L1 CSM responsive neurons, we aligned the data to L1 protrusion onset and assessed significance -100 ms to 0 ms relative to protrusion onset (pre-protrusion onset responsive) or 0 ms to 200 ms relative to L1 protrusion onset (post-protrusion onset responsive). To assess significance for firing rate differences in peri-L1 responsive neurons, and in all subsequent analysis, we performed a Wilcoxon-signed rank (WRS) test on the number of spikes in the two trial types under comparison in 25 ms windows. Windows were shifted in 5 ms steps and considered significantly modulated when at least 4 consecutive windows exhibited p < 0.05.

Neurons significantly modulated by L1 CSMs in any of the three epochs defined above were classified as ‘selective’ within their respective epoch, and further assessed for trial-type selectivity^31^. Peri-stimulus firing rate histograms (PSTHs) were constructed with 10 ms bins and smoothed with a 3-bin moving average. Selectivity at each time bin was defined as the absolute spike rate difference between trial types, normalized by peak selectivity. Selectivity heatmaps for each epoch (Fig. 4) were sorted in descending order by median selectivity within the epoch.

To test for neuronal correlates of double-step trials (Fig. 6a-b), we assessed the difference in firing rate between double-step and control trials. Data were aligned to L2 protrusion onset and tested for significance from 0 - 400 ms relative to L2 protrusion onset. To classify whether a neuron was activated or suppressed in response to double-step trials, we calculated the median difference in selectivity of each neuron within the epoch relative to control trials. Neurons were classified as activated if this difference was greater than 0, and suppressed if this difference was less than 0.

To test for neuronal correlates of premature bout termination (Fig. 6d), we quantified the difference in firing rate between trials where bouts were terminated and trials where bouts continued. A bout was considered to be terminated if there were no licks after L2, and considered to be continued if there was at least one lick after L2. Data were aligned to L2 protrusion onset, and tested for significance from 0 - 150 ms relative to L2 protrusion onset.

To test for neuronal correlates of L2 spout misses (Fig. 6e), we compared the difference in firing rate between trials where L2 made spout contact with trials where L2 did not make spout contact. To control for termination signals, we only included trials where there was at least one lick after L2. Data were aligned to L2 protrusion onset, and tested for significance from 0 - 150 ms relative to L2 protrusion onset.

To test for neuronal correlates of L2 spout position (Fig. 6g), we assessed the difference in firing rate between trials where L2 made spout contact on double-step (spout far) relative to control (spout near) trials. To control for termination signals, we only included trials where there was at least one lick after L2. Data were aligned to L2 spout contact onset, and tested for significance from 0 - 100 ms relative to L2 spout contact onset. For clarity and consistency, panels in Fig. 6 were plotted aligned to L2 protrusion onset.

Finally, to test for neuronal correlates of L2 CSMs (Fig. 6f), we compared the difference in firing rate between trials where L2 contained CSMs versus trials where L2 lacked CSMs. To control for termination signals, we only included trials where there was at least one lick after L2. Additionally, to control for spout contact signals and spout contact position signals, we only included double-step trials where L2 missed the spout. Data were aligned to L2 protrusion onset, and tested for significance from -50 - 100 ms relative to L2 protrusion onset. All analyses for L2 described above were repeated for L3 (Extended Data Fig. 11).

To determine when ALM activity on double-step trials diverged from control trials (Fig. 6c), we first used principal components analysis to reduce the dimensionality of our data as described previously^9, 10, 32^. As dimensionality reduction methods can be biased by high firing rate units, we normalized each neuron’s firing rate by the maximum standard deviation for each unit across all trials and all conditions^9, 10^. Data were aligned to L2 protrusion onset, and PSTHs were generated in 10 ms bins as described above. We then ran PCA on this data and projected the condition-averaged (double-step, control) responses onto the top 8 dimensions of this space, which explained >95% of the neural variance in our dataset. We plotted the trajectories from each condition in the first three dimensions of this space or trajectories in two of the top three dimensions in this space (Fig. 6c). To estimate the neural distance between trajectories and variability of this distance, we performed a bootstrap analysis. For each condition (double-step, control), we resampled trials with replacement for that condition of equal size as the original dataset. We then computed PSTHs with this resample dataset, projected the data onto the top 8 principal components, and calculated the euclidean distance between these trajectories. This procedure was repeated 1000 times to yield the bootstrapped mean neural distance between trajectories and an estimate of the variability in the distance between neural trajectories.

### Electrophysiological validation of Photoinhibition

To validate the photoinhibition (Extended Data Fig. 7), we performed acute extracellular neural recordings in awake VGAT-Chr2-EYFP animals (2 animals, 2 sessions) while simultaneously performing photoinhibition with laser powers identical to those used in the behavioral tests (40Hz sinusoidal wave at 10mW). Photoinhibition was delivered for 1.1 seconds with an exponentially distributed time interval (rate parameter: 3 seconds) between inactivations.

### Statistical analyses of tongue kinematics

Statistical analyses were performed using standard tests in MATLAB, including one sided t-tests, two-sample t-tests, Wilcoxon rank sum tests and Wilcoxon signed-rank tests. Correlation was tested by applying the F-test statistic to a linear fit. A chi-squared goodness-of-fit test was performed to determine if a distribution was uniform. For measures of central tendency, we used medians and interquartile ranges (IQR) since these measures do not assume normality of distributions. They are represented as medians [IQR]. For e.g. a duration measure of 18ms [16 22], represents a median of 18ms with interquartile range from 16ms to 22ms. For all estimates of kinematic parameters, animals were only included if they had at least 5 data points in each condition.

We generated linear mixed effects models to test if the probability (CSM_pres) or the duration of the CSMs (CSM_dur) in the first cue-evoked lick could be predicted by time since last spout contact (prev_spc), trial number in the session (trial_num) or the reaction time (RT). (Extended dat fig 6c-d). For modelling the probability of CSMs we used the formulation: CSM_present ∼ 1+ prev_spc + trial_num + RT + (1|MouseID) + (RT|MouseID), with CSM_present as a binomial distribution and a logit link function. For modelling the duration of CSMs we used the formulation: CSM_durations ∼ 1 + prev_spc + trial_num + RT + (1|MouseID) + (RT|MouseID) with CSM_durations as a normal distribution and an identity link function.

## Supporting information

Movie S1 Example Cue-evoked and Retrieval licks

Movie S2 SpoutContact_AltersLicks

Movie S3 Example ALM Inactivation

Movie S4 Example Cue-evoked and retrieval licks (spout left)

Movie S5 ALM Inactivation (Spout Left)

Movie S6 ALM inactivation on L1

Movie S7 Double-step experiment

Movie S8 Double-step experiment with ALM inactivation

## Acknowledgments

We thank Joe Fetcho, Melissa Warden, Neville Hogan and members of the Goldberg lab for comments on the manuscript. We thank Bilal Bari, Nikil Prasad and Jackson Walker for technical advice.

## Author contributions

TB, BI and JHG designed the research; TB and BI performed the experiments; TB, BI, SW, and JHG analyzed the data; BK, TB, and BI hardware; BI, ML and TB performed surgeries and histology; TB, BI, SW and JHG wrote the paper. The authors declare no competing financial interest.

**Extended Data Fig. 1.**
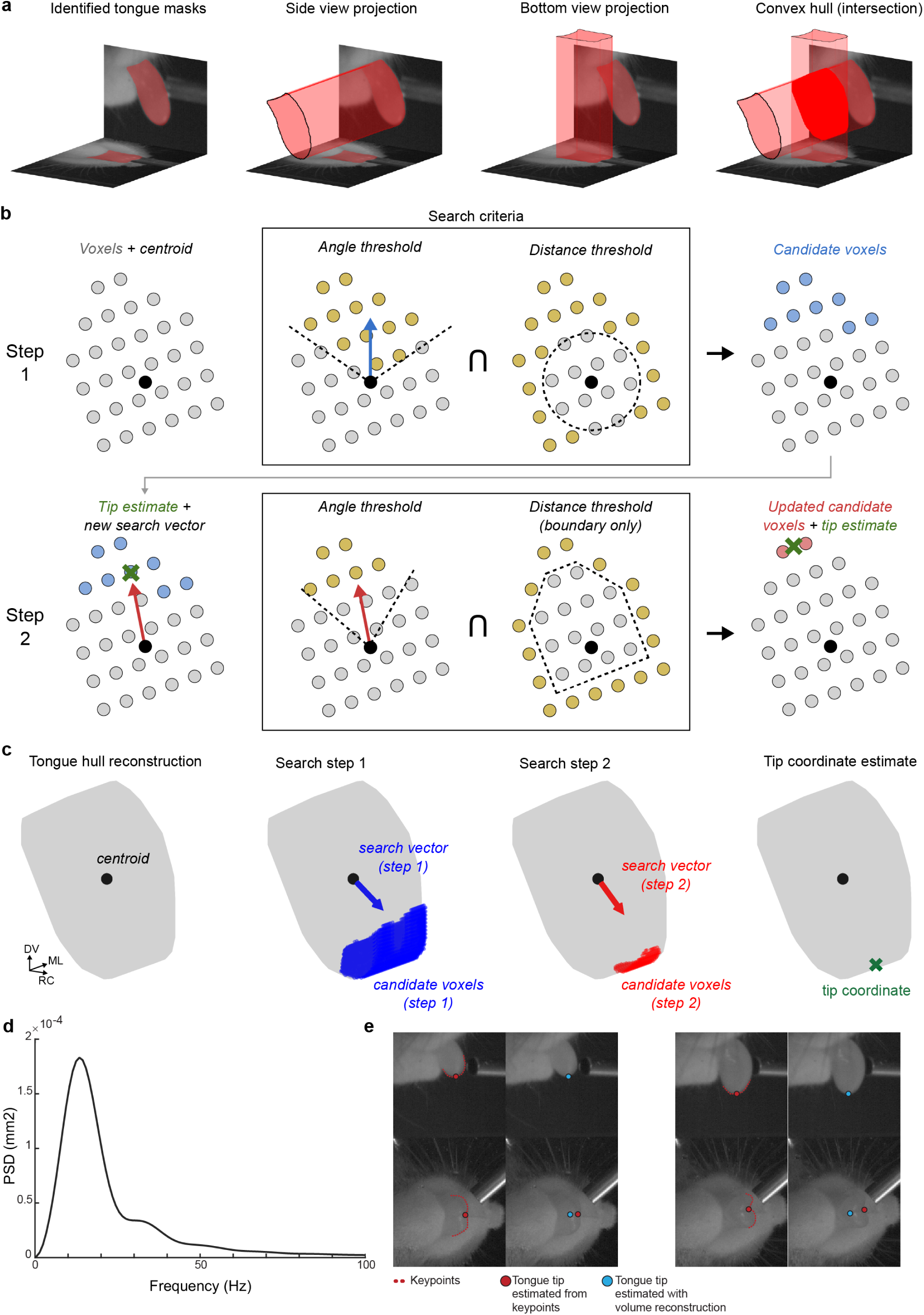
Method for extracting 3D tongue tip kinematics. **a**, An example of the process used to generate a 3D voxel hull from the two views of the mouse tongue. The walls of the diagrams are stills taken from the high-speed video, with the segmented tongue mask highlighted in red. The final hull (rightmost diagram) is obtained by identifying the intersection of the projections of the side and bottom view tongue masks. **b**, A 2D illustration of the tip coordinate search. With the voxels (gray circle) and centroid (black circle) identified, the first search step is performed, in which candidate voxels (blue) are found via the intersection of voxels satisfying the two search criteria (yellow), namely thresholds on the maximum angle made with the search vector (blue arrow) and the minimum distance from the tongue centroid. These first candidate voxels are then used to generate a refined search vector (red arrow, second row) for the second step of the search. Using this refined search vector, a similar set of angle and distance thresholds are applied, to determine a refined set of candidate voxels, which are then averaged to determine the tip location. **c**, Example of the tip search process with real data in 3D. The gray object is the 3D tongue hull, with the centroid labelled by a black circle. The first search step identifies a set of candidate voxels (blue) that are used to generate a refined search vector for the second search step (red). Using the second step candidate voxels, the tongue tip location is estimated (green x). **d**, Average power spectral density plot of tongue tip trajectories from five representative animals. >90% of power was at frequencies less than 50 Hz. **e**, Two time-points of a single lick (left and right) are shown with tongue tip estimated with keypoints (red) and with volume reconstruction (blue). Though keypoint tracking appears to work well from the side view, it fails in the bottom view since the true tongue tip does not always lie at the edge of the image of the tongue as seen from the bottom. This is because in most licks the tongue exhibits a ‘c’ shape at full extension. In these frames the tip is mislabelled by keypoints in the bottom view. Importantly the error cannot be accounted for systematically because it varies dynamically within a lick according to the tongue’s convexity.

**Extended Data Fig. 2.**
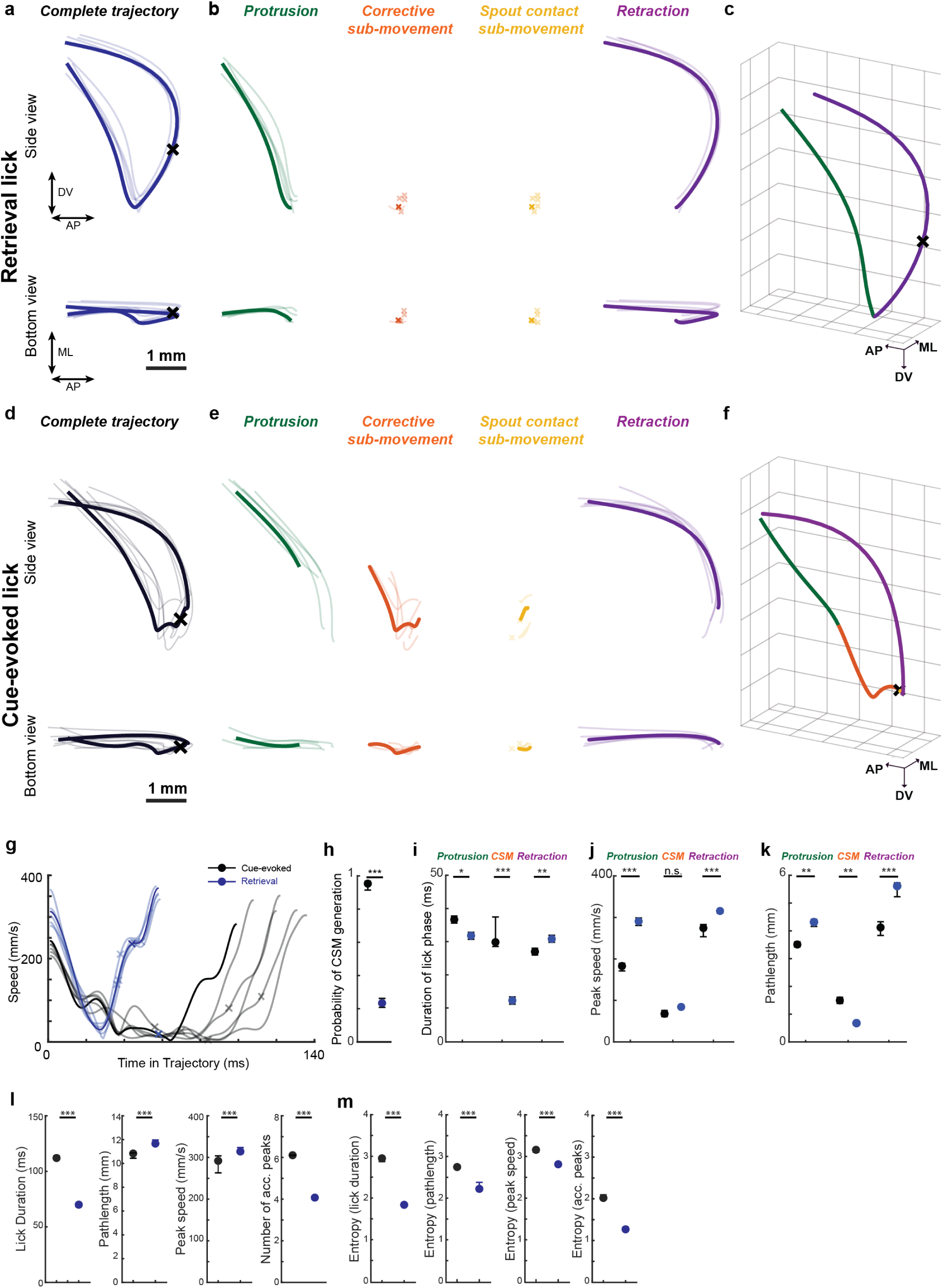
Water retrieval and cue-evoked licks exhibit distinct kinematics. **a-c**, Water retrieval licks, defined as those initiated after spout contact. **a**, Six overlaid tongue tip trajectories during retrieval licks. A single lick is bold for clarity. **b**, Protrusion, CSM, SSM, and retraction phases of the trajectories from **a** are separately plotted. x symbols denote the absence of CSMs and/or SSMs. **c**, 3-D trajectory of the highlighted lick shown in **a**, with protrusion (green) and retraction (purple) lick phases indicated. **d-f**, Data plotted as in **a-c** for cue-evoked licks. Note the prominent CSMs. **g**, Tongue tip speed profiles for retrieval (blue) and cue-evoked (black) trajectories shown in **a,d. h-k**, Probability of CSMs **(h)** and durations **(i)**, peak speeds **(j)** and pathlengths **(k)** of distinct lick phases during cue-evoked (black) and retrieval (blue) licks. **l-m**, Kinematics **(l)** and Entropy **(m)** of lick durations, pathlengths, peak speeds, and number of acceleration peaks. Data in **h-m** are median±IQR across 17 animals all from sessions with spout at 3.2mm.*, ** and ** denote p<0.05, <0.01 and <0.001 for a wilcoxon signed rank test.

**Extended Data Fig. 3.**
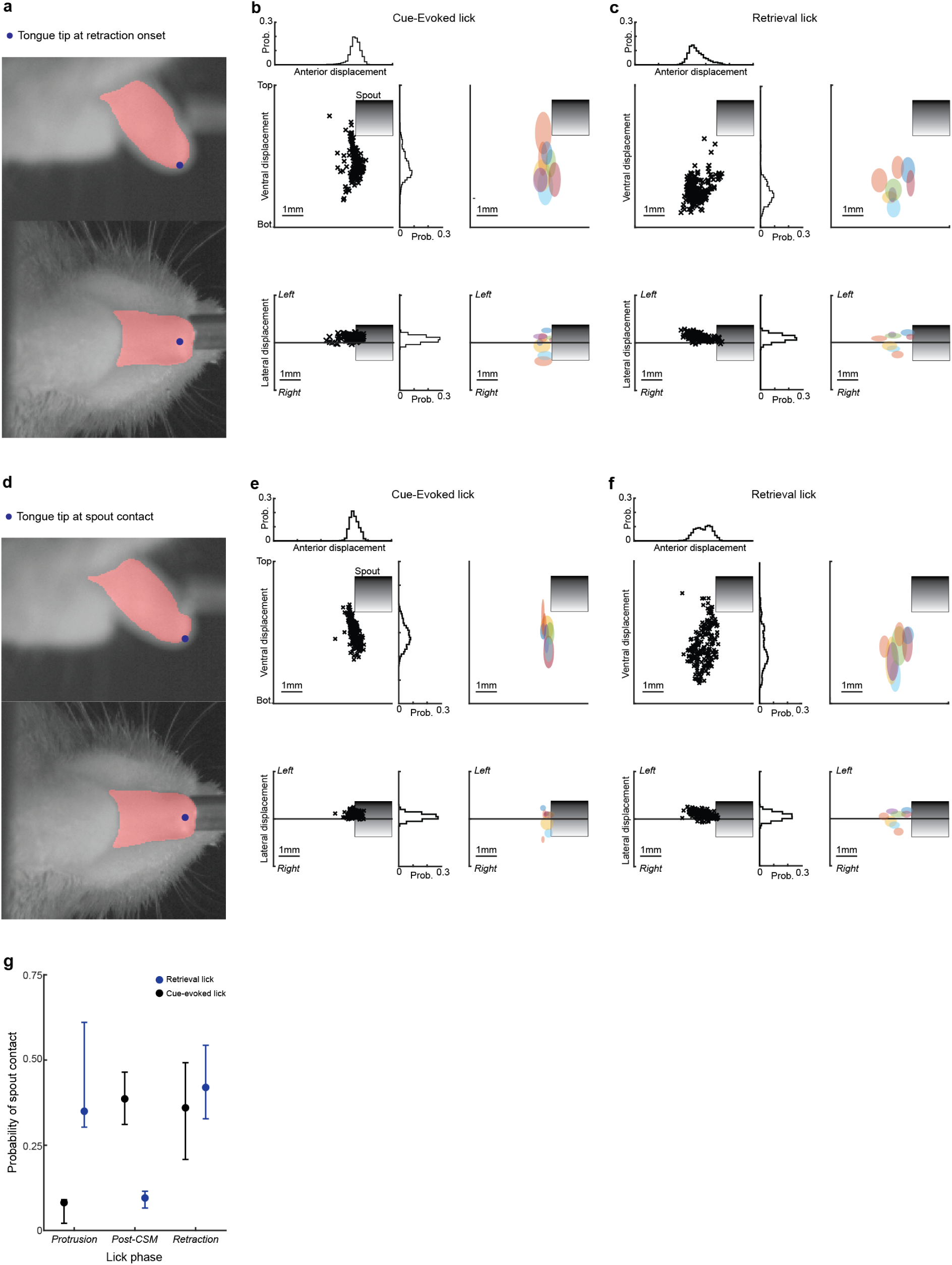
Tongue tip positions at the moment of retraction onset and spout contact. **a**, Side and bottom views of the tongue at the moment of retraction onset from a representative lick. **b**, Scatter plots of tongue tip positions at retraction onset for side (top) and bottom views (bottom) during successful cue-evoked licks from a single session?. Probability distributions are projected along the axes at top and right (bin size, 120um). Right, 2D standard deviations of tongue tip positions at retraction onset for 9 representative mice (each mouse independently color-coded). Note that each mouse exhibits a ‘preferred’ target location for retraction onset. **c**, Tongue tip positions at moment of retraction onset plotted as in b for retrieval licks. **d-f**, Data plotted as in **a-c** for tongue tip positions at the moment of spout contact (n=the same 9 mice). **g**, Probability of spout contact as a function of the distinct lick phases for cue evoked and water retrieval licks (blue and black, respectively, n=17 mice).

**Extended Data fig. 4.**
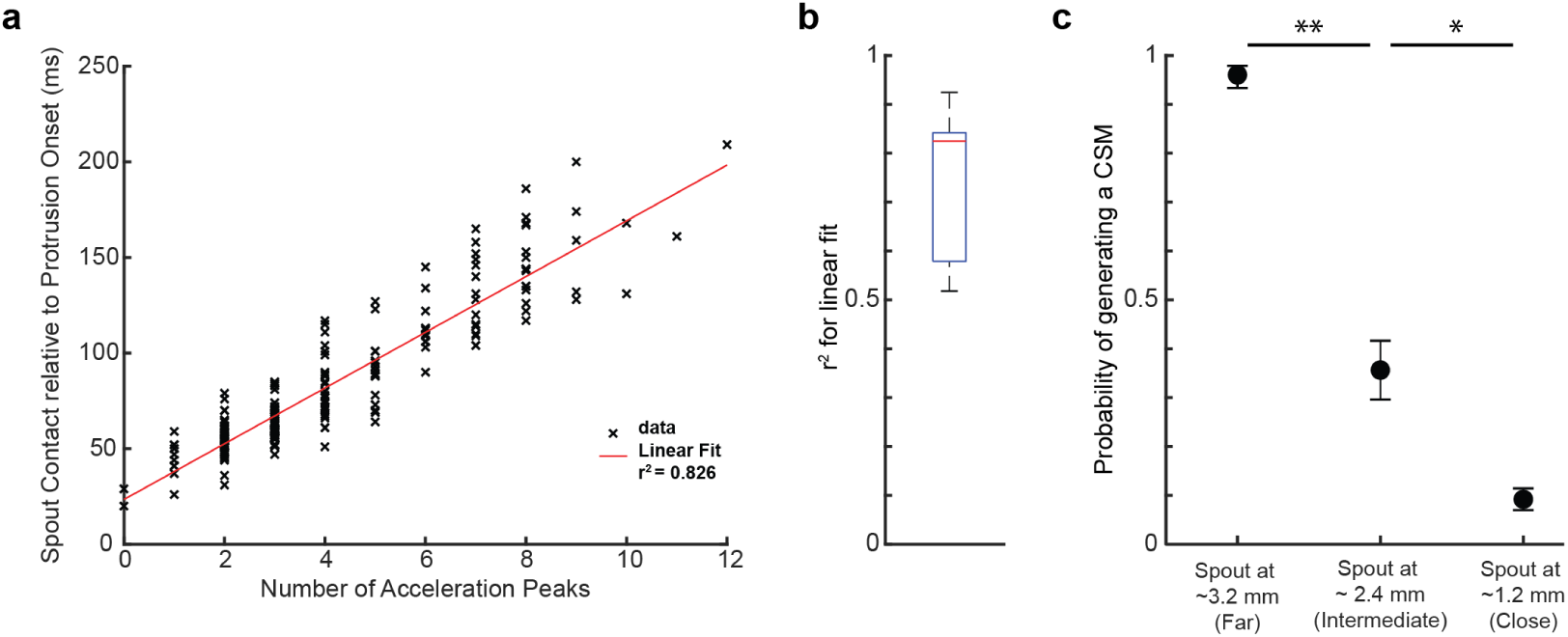
The number of acceleration peaks per lick predicts latency to spout contact. **a**, The latency to spout contact relative to protrusion onset is plotted against the number of acceleration peaks per lick from a single spout-far session. Red line, linear fit. **b**, Boxplot showing r^2^ for linear fits across 17 animals (red line: median; box edges: IQR; whiskers: 95% CI). c, CSM probability scales with spout distance (n=17, 11, and 13 animals for spout at 3.2, 2.4 and 1.2 mm respectively;*,** denote p<0.05, <0.01 for a wilcoxon rank sum test)

**Extended Data fig. 5.**
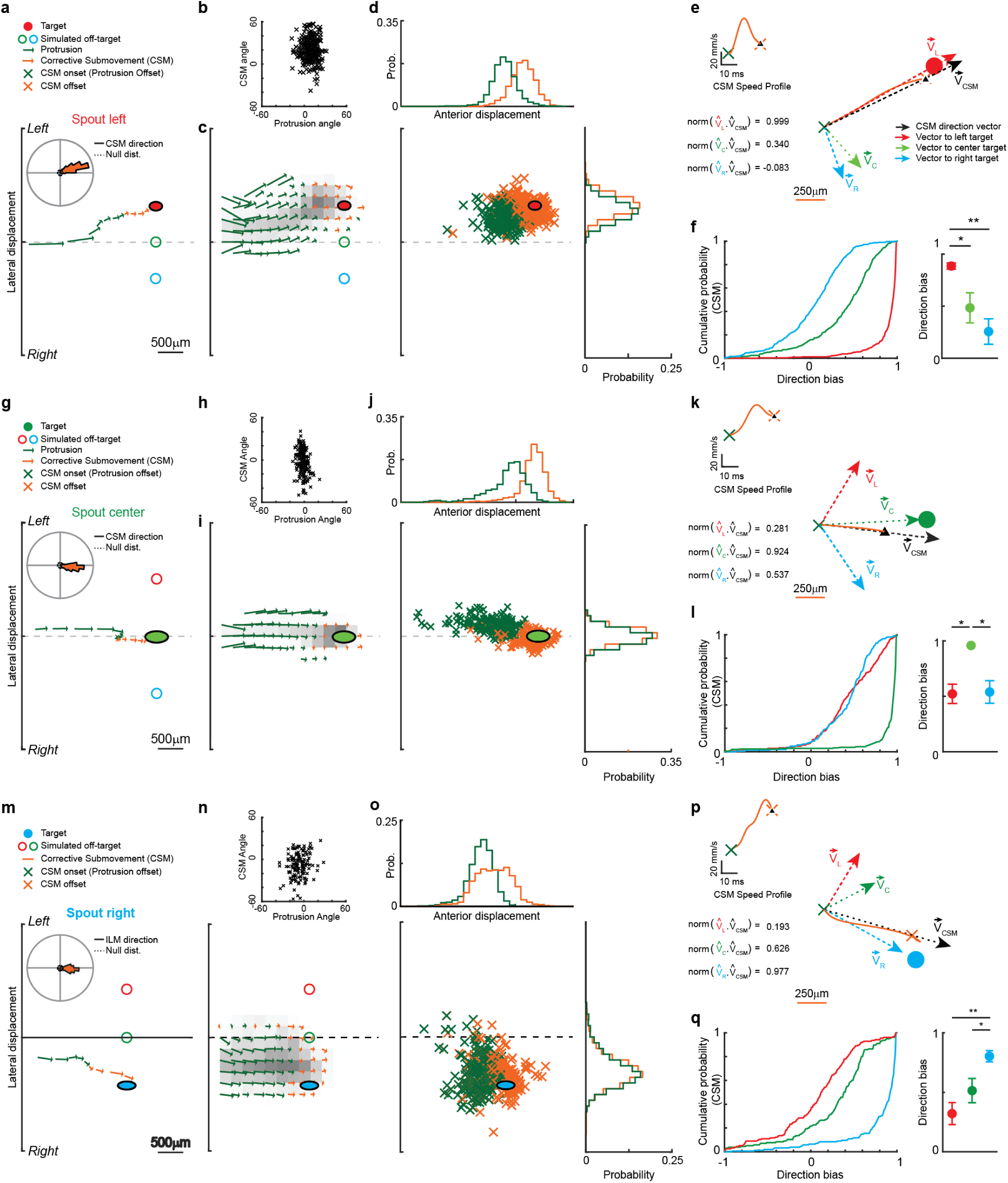
Corrective submovements are directionally biased towards remembered spout locations. **a-e**, CSM kinematics for spout-left sessions. **a**, time-dependent velocity vector for the protrusion (green) and CSM (orange) phase of a single cue-evoked lick. The origin of each vector is the tongue tip position at 5 millisecond intervals of the lick, the amplitude is the speed, and the arrow points in the direction. Inset: polar plot with direction distribution of all CSMs produced in a single spout-left session. Dashed circle, the null distribution of unbiased CSM directions. **b**, Scatter plot of CSM directions plotted against protrusion directions for all cue-evoked licks in the session. **c**, Position-dependent average velocity vectors for all cue-evoked licks from a single session, color coded by highest likelihood lick phase to pass through the binned space (250um grid). Gray shading intensity of each bin is proportional to the probability of a tongue tip trajectory passing through the space. **d**, Scatter plot of tongue tip positions at protrusion offset (green) and retraction onset (orange), indicating CSM start and end points. Probability distributions of the CSM start- and end-points are projected along the axes at top and right (bin size, 120um). **e**, Example of a single CSM path and its speed profile (orange). The CSM’s initial direction (Vilm, black dotted line) was computed from the vector connecting CSM starting point to its position at the first speed minimum (upward black triangle in speed and path plots). The dot products between this CSM direction vector and three additional vectors from CSM starting point to left, center, and right targets (dotted red, green, and blue lines, respectively) were computed to quantify the ‘direction bias’, the extent to which the initial direction of a given CSM was aimed at each of the three candidate targets. **f** Left: Cumulative distributions of directional biases for all CSMs produced in a single session to the three candidate targets (colored as in **b**). CSMs were reliably aimed to the left target. Right: Directional biases of CSMs to left, center, and right targets spout-left sessions (median±IQRs across n=13 animals,*, ** denote p<0.05, <0.01 for Wilcoxson ranked sum tests. **(g-l)** CSM kinematics for spout-center sessions, plotted as in **a-f** (n = 17 animals). **(m-q)** CSM kinematics for spout-right sessions, plotted as in **a-f** (n = 12 animals).

**Extended Data fig. 6.**
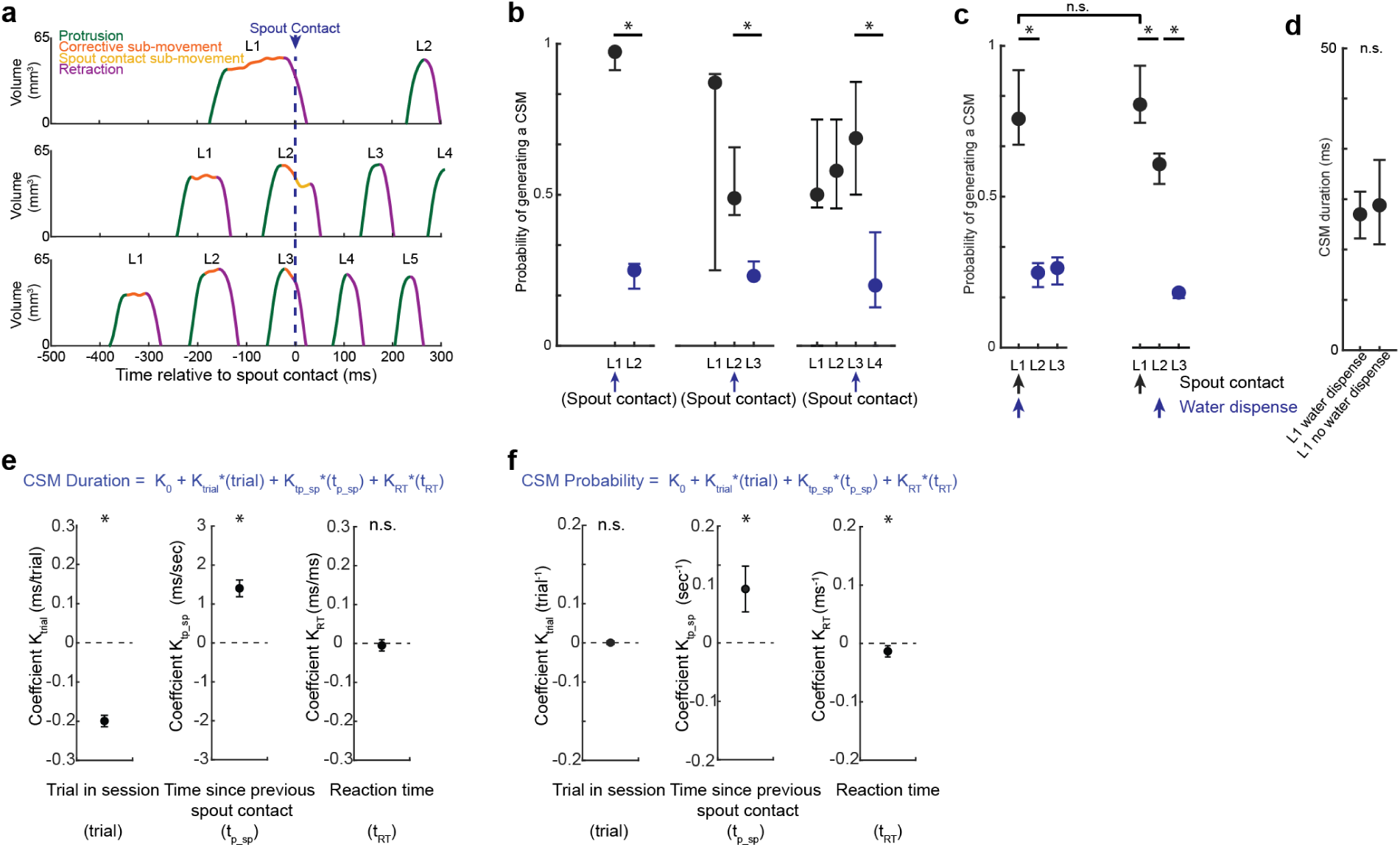
The first spout contact transforms the kinematics of subsequent licks in a bout. **a**, Tongue volumes as a function of time during three trials where first spout contact (black dashed line) occurred on the first, second, or third lick. Note that licks initiated before spout contact exhibited substantial CSMs, whereas those initiated after spout contact did not. **b**, CSM probability as a function of lick number in cases where first spout contact happened on first, second, or third licks (median ± IQR across animals n=17 animals). Spout contact reliably transformed the kinematics of subsequently initiated licks. Data in **b** from sessions where water was dispensed on first spout contact. **c-d** CSM probabilities when water was dispensed on the second spout contact **c**, CSM duration on first cue-evoked licks (L1) did not depend on water dispensation on first contact. **d**, In sessions where water was dispensed on second contact, both spout contact on L1 and water dispense on L2 contact reduced CSMs on ensuing licks (n= 12 mice, *, p<0.05 wilcoxon signed rank test). **e-f**, Mixed effects models were used to predict the duration **(e)** and probability **(f)** of CSMs on first licks of a bout (Methods). CSM durations were significantly predicted by trial number in session and time since previous spout contact (t_p_sp_), but not reaction time (tRT). CSM probabilities were predicted by t_p_sp_ and by t_RT_.

**Extended Data fig. 7.**
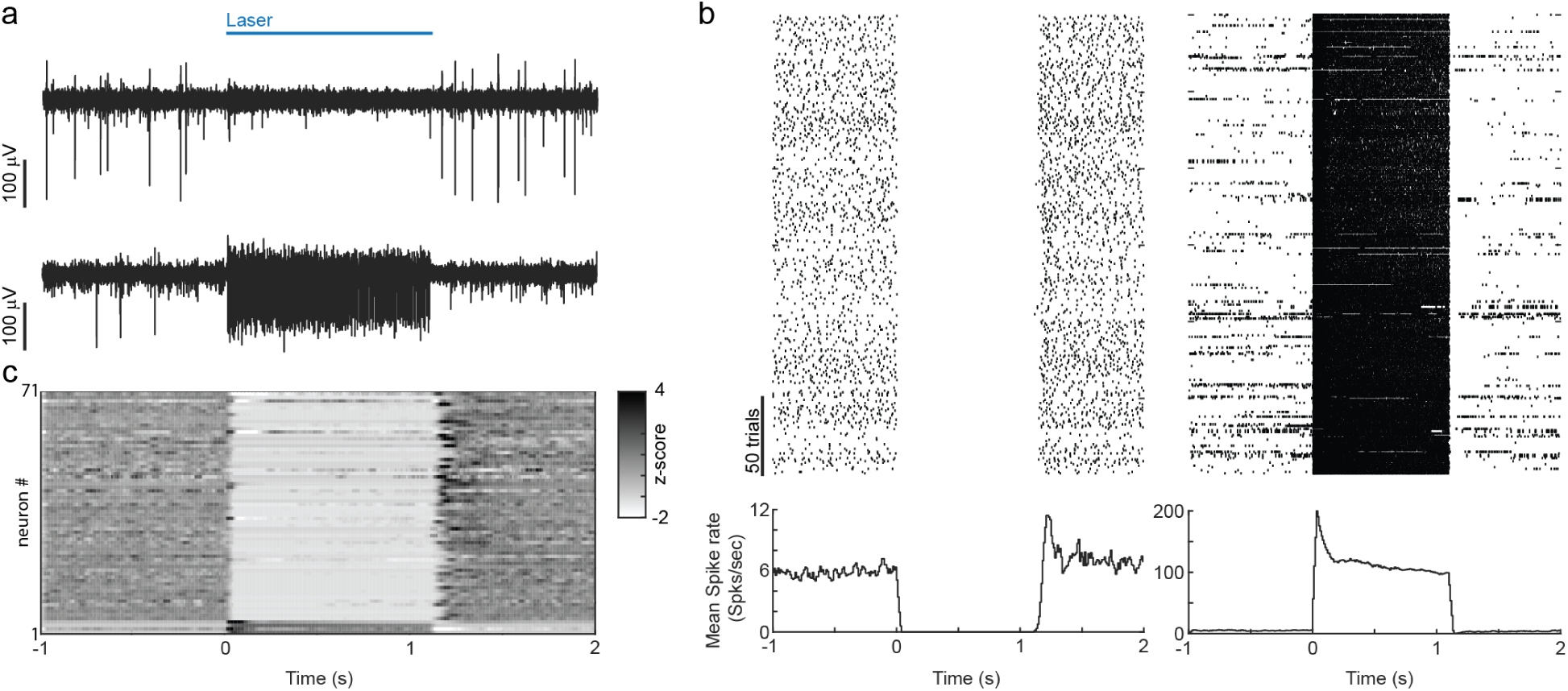
Electrophysiological validation of photoinhibition in VGAT-ChR2-EYFP mice. **a**, Voltage waveforms of putative pyramidal neuron (top) and interneuron (bottom) during one second illumination of 40Hz sinusoidal wave at 10mW, the same power and waveform generated in behavioral experiments. **b**, spike rasters and corresponding rate histograms of the neurons from (a). **c**, z-scored firing rates of 71 ALM neurons before, during and after optogenetic activation of inhibitory interneurons in VGAT-ChR2-EYFP mice (n=2 sessions, n=2 mice, Methods).

**Extended Data fig. 8.**
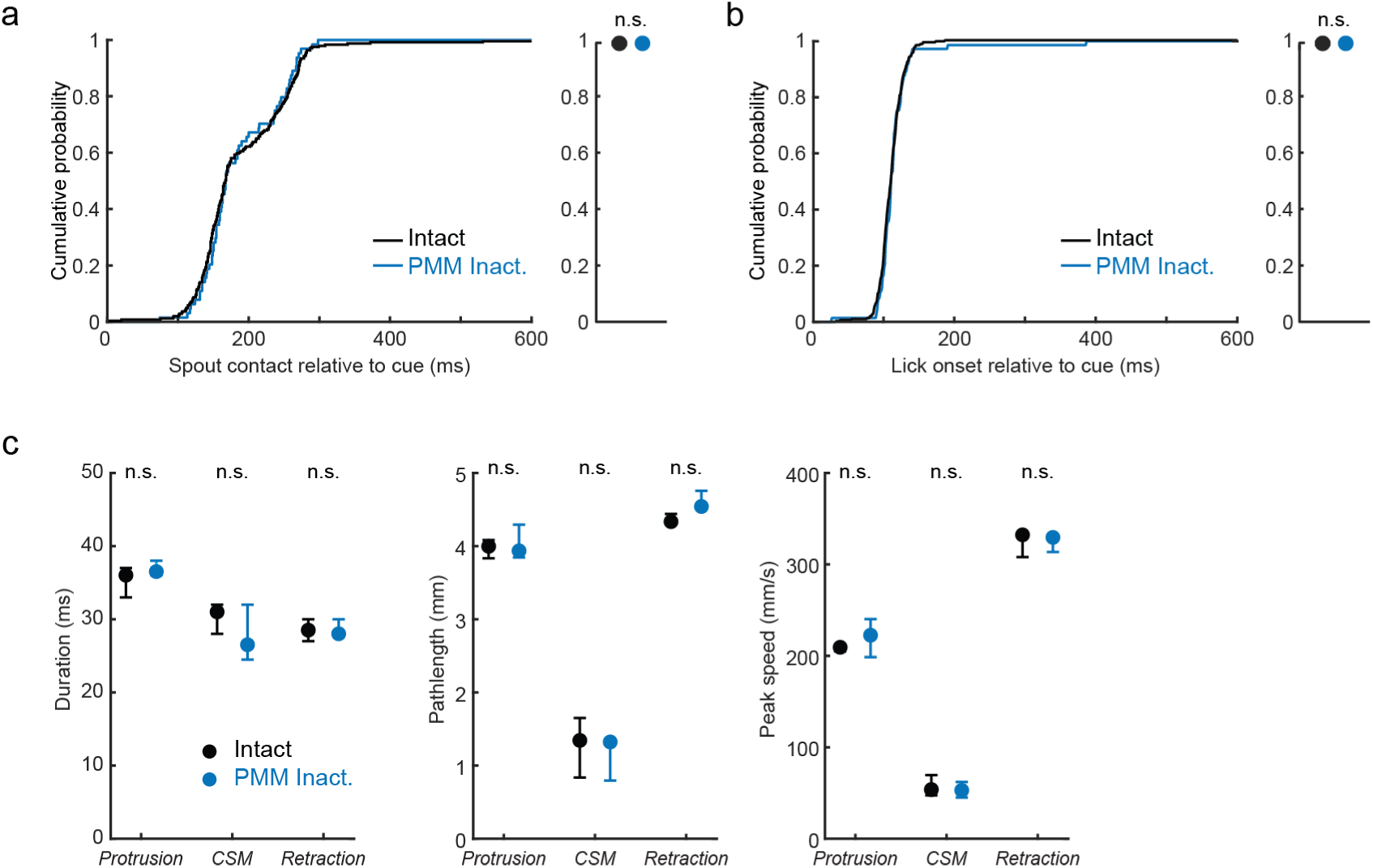
Inactivation of PMM does not impact task performance or lick kinematics. **a**, Cumulative probability of tongue-spout contact relative to cue onset during laser-off and PMM-photoinactivated trials. Right, median ± IQR probability of spout contact within a trial across mice (n = 9 mice). **b**, Data plotted as in **a** for tongue protrusions. **c**, Median durations, pathlengths, and peak speeds for lick phases with PMM intact (black) and PMM inactivated (blue), (median±IQRs).

**Extended Data fig. 9.**
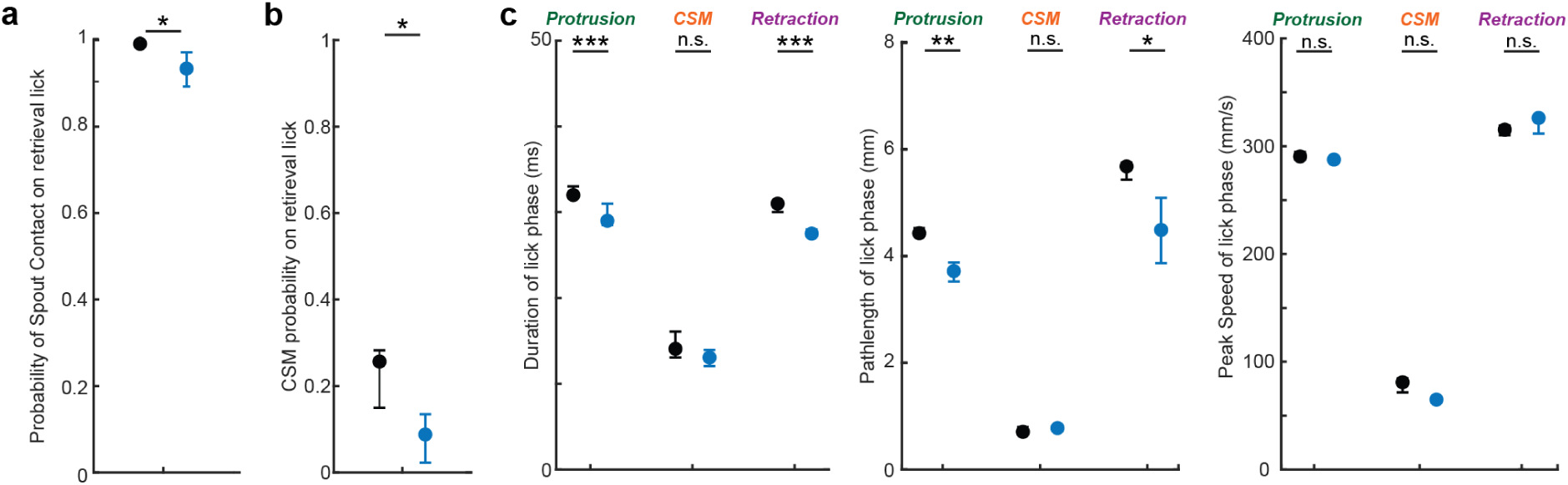
Impact of ALM photoinhibition on water retrieval licks. **a-b**, Spout contact **(a)** and CSM **(b)** probabilities were reduced by ALM photoinhibition (blue; control trials: blsck). **c**, Median durations, pathlengths, and peak speeds of retrieval lick phases produced with ALM intact and inactivated. Data are median ± IQRs of the first retrieval lick that followed cue-evoked licks that made contact during ALM photoinhibition (*, **, ***, denote p<0.05,0.01 and 0.001 in a Wilcoxon signed rank test; n = 12 mice).

**Extended Data fig. 10.**
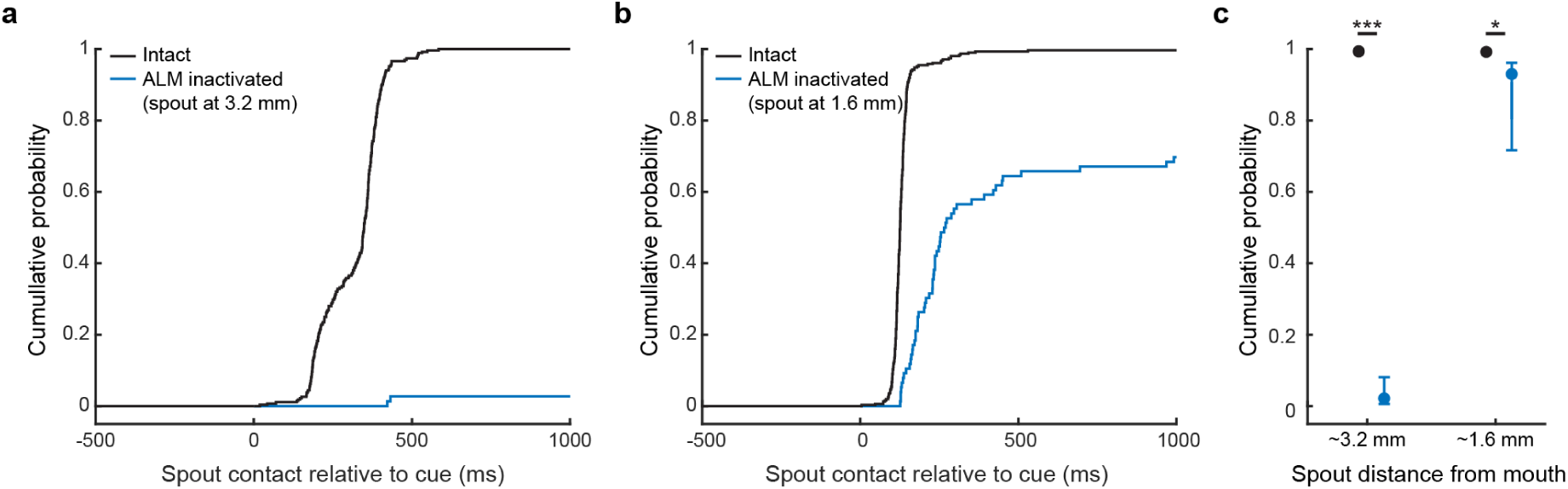
Proximal spout placement rescues ALM inactivation-associated spout contact deficits. **a-b**, Cumulative probability of spout contact relative to cue onset for control (black) and ALM-inactivated (blue) trials in sessions where the spout was 1.6 mm **(a)** and 3.2 mm **(b)** from the incisors. **c**, Median probability of spout contact across animals from spout-close and spout-far sessions (* denote p<0.05 for a Wilcoxson signed rank test n=13 animals).

**Extended Data fig. 11.**
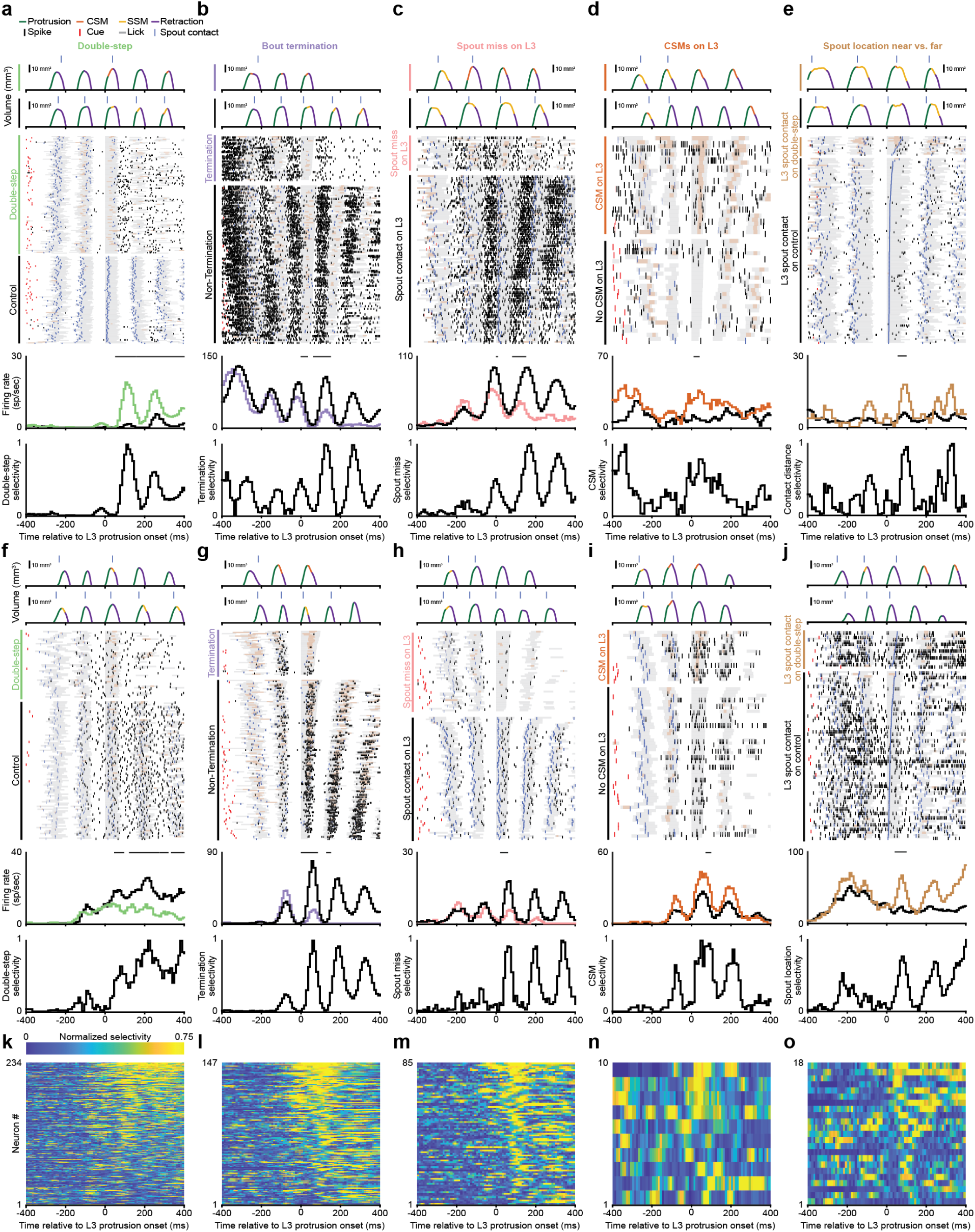
Neural correlates of online corrections on the third lick (L3) of double-step trials. **a**, Tongue volumes, spike rasters, and corresponding rate and double-step-selectivity histograms for 2 example ALM neurons. Neural activity aligned to L3 protrusion onset. Raster color-codes as in Fig. 4. Bottom, ALM population double-step selectivity, defined as the normalized difference in firing rate from control and double-step trials (Methods). Only neurons with significant trial selectivity are shown (n=234/465 neurons). **b-e**, Data plotted as in a for the following conditions: premature bout termination following L3 (n=147/438 neurons) **(b)**, L3 spout misses resulting in bout continuation (n=85/418 neurons) (c), CSMs on L3 misses (n=10/79 neurons) **(d)**, and spout location on L3 contacts (n=18/167 neurons) **(e)**.

